# Saturation barcoding of ribosomal DNA resolves the nucleolar organizer regions of the human genome

**DOI:** 10.64898/2026.11.10.765783

**Authors:** Shudi Zhao, Jakob Wallentin, Tamara Potapova, Shengzhang Zheng, Xinrui Liu, Riza M. Daza, Jennifer L. Gerton, Jay Shendure, Xiaoyi Li

**Author notes:** These authors contributed equally (Shudi Zhao, Jakob Wallentin).

## Abstract

The nucleolus is the hub for ribosome biogenesis in eukaryotic cells. It is assembled on nucleolar organizer regions (NORs), tandem arrays of ribosomal DNA (rDNA) units. In humans, each rDNA unit is ∼45 kb and encodes the 47S ribosomal RNA (rRNA) precursor. Hundreds of rDNA units are present per germline genome, spread across 10 NORs positioned on the short arms of the five acrocentric chromosomes. However, their precise count and structure are unknown. In fact, twenty-five years after the first draft of the human genome, NORs are the <u>only</u> regions of the human genome that remain incompletely resolved, represented in telomere-to-telomere (T2T) reference genomes as gaps or modeled sequences rather than bona fide assemblies. Here we use prime editing to insert degenerate 6-bp DNA barcodes to 99% of taggable rDNA units in a clonal derivative of the leukemia-derived K562 cell line. Following ultra-long-read sequencing, these barcodes serve as molecular identifiers that guide the end-to-end assembly of all 18 NORs of K562’s pseudotriploid genome. The assembly yields: (i) a precise rDNA copy number for each NOR (range 5-43; total 290); (ii) the *cis-* and *trans-*architecture of intragenomic variation in rDNA sequence, both structural and fine-scale; and (iii) patterns of divergence and homogenization within and across NORs that may have arisen during cancer evolution and the subsequent passaging of this line. Finally, by reading the same tags from RNA, we define the relationship between DNA methylation and Pol I transcription at the resolution of individual rDNA units. Together, our study demonstrates how the final tangles of the human genome can at last be resolved, and opens the door to more extensive investigations of how genetic, epigenetic and functional rDNA variation contributes to health and disease.

## INTRODUCTION

Ribosome biogenesis lies at the heart of the central dogma: it is the ribosome that translates genetic information from mRNA into protein. NORs are the genetic and structural foundation of this process, encoding the ribosomal RNA (rRNA) precursor (47S in humans) and seeding formation of the nucleolus itself (*1*, *2*). Yet ∼70 years after the central dogma’s first articulation, and ∼25 years after the first draft of the human genome, human NORs remain a blind spot in three respects. <u>First</u>, their reference architecture has never been fully resolved, stymieing each successive generation of sequencing technology. <u>Second</u>, variation in rDNA units is poorly characterized, whether within one genome, across human populations, or in cancer evolution. <u>Third</u>, how NOR architecture and variation shape ribosome function and human phenotypes remain underexplored.

The germline human genome contains 10 NORs, located on the short arms of the five acrocentric chromosomes (chrs. 13-15, 21-22). Each NOR is a tandem array of ribosomal DNA (rDNA) units. Each ∼45 kb rDNA unit comprises a ∼13 kb region encoding the 47S pre-rRNA, cotranscriptionally processed into the 18S, 5.8S and 28S rRNAs, and a ∼30 kb intergenic spacer (IGS) that is itself internally repetitive, harboring a spacer promoter, a Pol I enhancer, a Pol I promoter, Sal box termination signals within R repeats, and tandem Butterfly/Long repeats (*3*).

Although individual rDNA units can be cloned and sequenced (*4*, *5*), the assembly of NORs is an outstanding technical challenge. Both the initial and finished versions of the human reference genome, assembled from paired Sanger reads of several hundred basepairs, represented the acrocentric short arms as multi-megabase stretches of “N”s (*6*, *7*). In the decade that followed, massively parallel sequencing transformed most other areas of genomics, but given their even shorter read lengths, disambiguating repetitive regions became if anything more difficult (*8*).

In the past few years, the long reads generated by single molecule sequencing technologies (*9*) have at last enabled “essentially complete” human genomes, *e.g.* the homozygous CHM13 cell line (T2T-CHM13) in 2022 (*10*) and the HG002 diploid benchmark in 2026 (*11*). However, their completeness is qualified because the NORs remain ambiguous. In T2T-CHM13, the five rDNA arrays are the only unresolved sequences, handled through a bespoke strategy of nanopore walks through sparse de Bruijn graphs, segmentation and clustering of units into “morphs” by sequence similarity, and copy number inference for each morph (*10*). All five NORs remain flagged as model sequences; two comprise a single primary morph tiled head-to-tail, while three are mosaics of interspersed morphs whose ordering could not be disambiguated. In HG002, nine of ten arrays are represented as gaps, sized by FISH (*11*). To our knowledge, only two human NORs have been confidently assembled at single-unit resolution: a 6-unit paternal chr13 NOR (*11*) and a 33-unit paternal chr14 NOR (*12*).

Apart from reference architecture, how rDNA copy number varies across germline genomes has been of longstanding interest (*13*). Early estimates based on short-read sequencing suggested striking diversity (*14*) but these estimates were confounded by batch effects (*15–17*). Orthogonal approaches now place total content at ∼400-600 units per diploid genome, apportioned very unevenly across chromosomes (from 0-2 to 150+ units per NOR haplotype) and with limited correlation between homologs (*18*). These findings are consistent with early studies that traced rDNA “electrophoretic karyotypes” through pedigrees (*13*). The numbers and distribution of rDNA units may serve as a failsafe against their loss driven by the unequal crossover and gene conversion that continually reshuffle these arrays, homogenizing units in *cis* and redistributing variants across chromosomes (*19–22*). Somatically, 47S rDNA copy number is frequently reduced in cancers despite their elevated ribosome biogenesis (*23–25*).

To what extent does the primary sequence of rDNA units vary? Ribosome heterogeneity arising from protein components has been studied extensively (*26–28*), but the genetic heterogeneity of its RNA components has received far less attention, presumably because such variation is difficult to survey. Variant-specific probes first revealed 28S rRNA heterogeneity within and between individuals, as well as tissue-specific expression of distinct variants (*29–31*). Sequencing subsequently resolved extensive heterogeneity across the rDNA unit, in both the rRNA-coding region and the IGS, with specific variants detected in translating ribosomes (*32–34*). Genetic associations have been suggested between rDNA variation and a range of human phenotypes, including gene expression, mitochondrial abundance, hematological traits, renal function, adiposity and body dimensions (*14*, *32*, *35*, *36*), although none are confirmed.

Despite 80-90% of cellular RNA is ribosomal, roughly half of rDNA units are silenced at any given time, with DNA methylation and histone modifications jointly specifying transcriptional state (*2*). Altered rDNA methylation accompanies aging, Down syndrome and neurodegeneration (*37–40*). However, absent a unit-resolved map, it has been difficult to determine which copies are transcribed, whether particular variants are preferentially silenced, and which specific NORs or rDNA units might underlie the suggested phenotypic associations.

Genome assembly, human genetics, and functional studies would all benefit from the ability to track individual rDNA units. As naturally occurring variation in these regions is sparse, we wondered whether we could instead introduce synthetic variation that facilitated such tracking. Synthetic barcodes have previously been used for genome and haplotype assembly, but usually as *in vitro* tags on molecules rather than marks written into the genome itself (*41–44*). For our objective here, we imagined that a diverse barcode pool, installed at a fixed position in every rDNA unit in the genome of living cells, would turn NOR assembly into a tractable ordering problem, while also enabling individual units to be assessed for sequence variation, methylation and transcriptional activity.

Conventional Cas9 editing is poorly suited for this rDNA barcoding objective, as double-strand breaks are cytotoxic in large numbers and inhibit Pol I transcription (*45*, *46*). In contrast, prime editing avoids double-strand breaks, specifies its outcome entirely within the pegRNA (*47*, *48*), is most efficient at transcriptionally active loci (*49*), and enables insertional targeting of repetitive sites at scale (*50*).

Here we show that prime editing of rDNA is highly efficient, and exploit this to install degenerate 6-bp barcodes into 99% of the taggable rDNA units of a clonal derivative of K562, a BCR-ABL-driven chronic myelogenous leukemia line bearing the Philadelphia chromosome, t(9;22)(q34;q11) (*51*, *52*). We then apply nanopore sequencing to this barcoded genome and, from the resulting reads, assemble all of its 18 NORs, end-to-end and anchored to a genome-wide assembly. The assembly yields the exact number of rDNA units in each NOR, the *cis-* and *trans*-architecture of intragenomic sequence variation, both structural and fine-scale, and patterns of divergence and homogenization within and across NORs that may have arisen during cancer evolution and the subsequent passaging of this line. Finally, by reading the same barcodes in RNA, we define the relationship between DNA methylation and Pol I transcription at single-unit resolution.

## RESULTS

### Multiplex prime editing of rDNA units is highly efficient

We previously found that prime editing efficiency is strongly correlated with the Pol II transcriptional activity of the target locus (*49*). Ribosomal DNA is transcribed by Pol I rather than Pol II, but at exceptionally high levels and within a distinct nuclear compartment. We wondered whether NORs were similarly permissive. Their repetitive architecture facilitates asking the question, as a single pegRNA species targets every rDNA unit (*50*).

We designed constructs encoding 11 pegRNAs to install 3-nt insertions at distinct positions across a chromosome 22 rDNA reference sequence (*5*) and transfected them into K562 or HEK293T cells stably expressing the prime editor PEmax (*49*, *53*). Ten of the 11 designs generated detectable edits. In K562 cells, eight exceeded 10% editing, and the most potent reached 62% (**fig. S1A**).

We narrowed our focus to K562 because of its higher prime editing rates (**fig. S1A**). To benchmark the rates observed in rDNA, we compared them against an active, well-characterized pegRNA targeting the HEK3 locus (HEK3.insCTT) (*48*). We selected 5 of the 11 rDNA pegRNAs, *i.e.* one (18S, ITS1, 28S) or two (IGS) per region (**Fig. 1A**), and transfected constructs encoding them into K562 cells alongside one encoding HEK3.insCTT. With HEK3.insCTT, we observed editing rates of 24%, 30% and 31% at 2, 4 and 6 days post-transfection, while with rDNA-targeting guides, we observed editing rates ranging from 5.4-36% at day 2, 11-54% at day 4, and 8.4-55% at day 6 (**Fig. 1B**).

**Figure 1.**
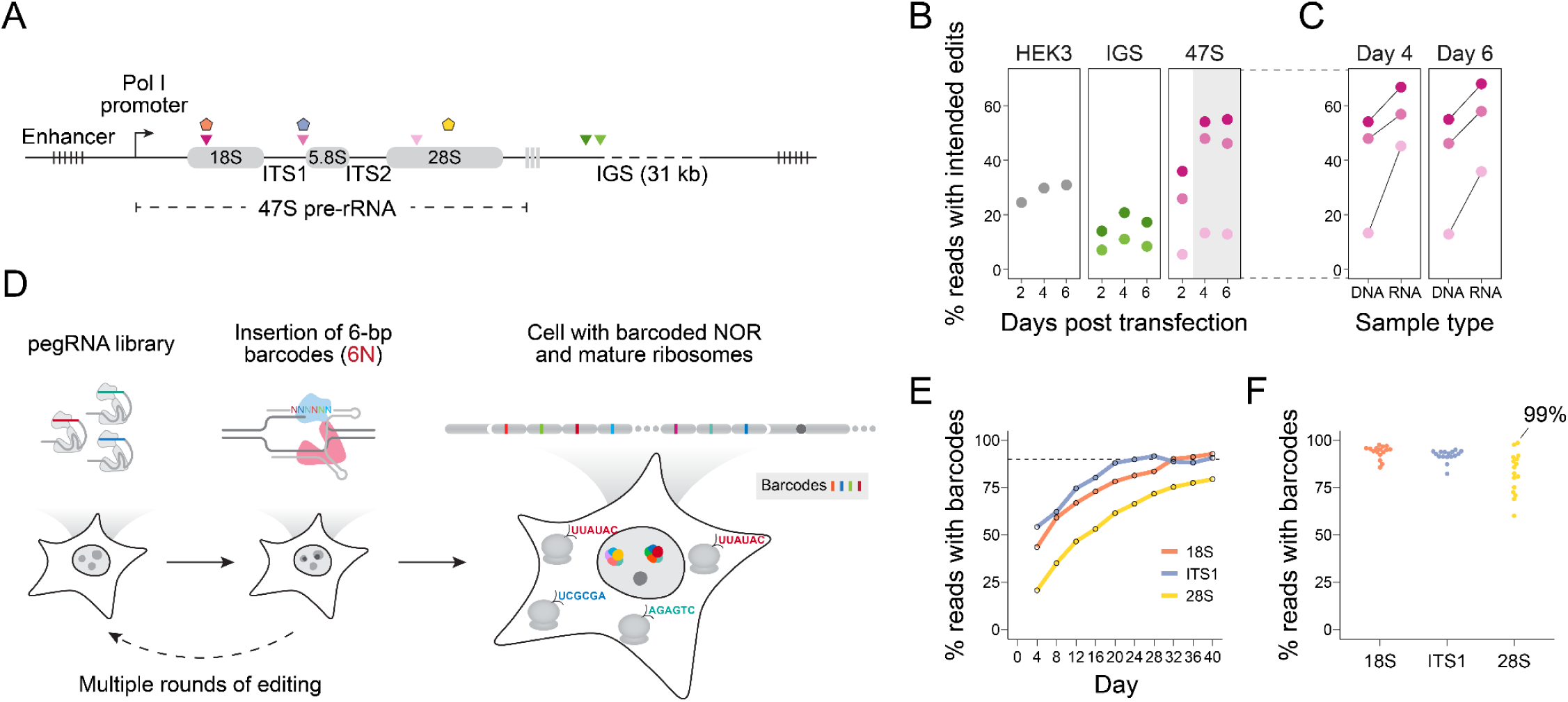
Saturation barcoding of rDNA units of the K562 genome by multiplex prime editing. **(A)** Schematic of an rDNA unit. Inverted triangles mark the target sites of the five pegRNAs used in **B-C**; the pentagons mark the barcoding sites used in **E-F**. ETS, external transcribed spacer; ITS, internal transcribed spacer; IGS, intergenic spacer. **(B)** Prime editing efficiency in gDNA at the *HEK3* locus, two IGS sites, and three 47S pre-rRNA sites in PEmax(+) K562 cells, 2, 4 and 6 days after transfection**. (C)** Editing efficiency of the three 47S-targeting pegRNAs in matched gDNA and rRNA from the same cells at days 4 and 6. **(D)** Schematic of the rDNA barcoding strategy. A pegRNA library installs 6-bp degenerate barcodes at a single fixed position in the rDNA unit; after repeated rounds of editing, monoclonal lines are derived so that barcodes map to individual rDNA units. **(E)** Percent of reads carrying a 6-bp barcode at the 18S, ITS1, and 28S sites over ten rounds of transfection in K562 PEmax(+) cells, measured every 4 days immediately before the next round. **(F)** Barcoding rates of the 16 monoclonal lines derived from each of the three pools in **E**. The 28S-barcoded clone used for NOR assembly is labeled with its barcoding rate of 99%,

We also asked whether the transcription levels of rDNA units are correlated with their editing rates, by measuring editing rates in rRNA from the same cells at the later time points. For every 47S-targeting pegRNA, editing rates were higher in rRNA than in matched genomic DNA (gDNA). The top-performing pegRNA, which targeted 18S, read out at 55% in gDNA and 68% in rRNA (**Fig. 1C**), suggesting that actively transcribed units are successfully edited at higher rates. This parallels our previous finding for Pol II-transcribed targets (*49*). Since the K562 cell line that we are using here carries ∼300 rDNA units (see below), a 55% gDNA editing rate corresponds to the simultaneous editing of ∼160 rDNA units per cell, on average.

### Saturation barcoding of the rDNA units of the K562 genome by multiplex prime editing

Having established that prime editing of NORs is highly efficient, we next asked whether it could be used to molecularly barcode individual rDNA units. Because prime editing writes an arbitrary sequence specified by the pegRNA, a library sharing a single spacer but differing in its programmed insertion can install a diverse set of barcodes at one fixed position across every unit in the genome (**Fig. 1D**). Specifically, we sought to introduce a 6-bp degenerate barcode: its 4⁶ = 4,096 possibilities exceed the expected number of rDNA units in the human genome by roughly an order of magnitude.

For this goal, the 18S and ITS1 sites from the benchmarking experiment (**Fig. 1A**) were the most efficient, but 18S is highly conserved, such that insertions into it risk disrupting ribosome assembly, while ITS1 is excised during rRNA processing and cannot report on mature ribosomes. We therefore designed a new pegRNA targeting 28S that coincides with an expansion segment that protrudes from the ribosomal surface (ES20L) (*54*) and is among the segments that varies naturally in human populations (*34*). Barcodes at ES20L are retained in mature rRNA, allowing ribosomes to be traced back to individual rDNA units (**Fig. 1D**; **fig. S1B**). We also confirmed that their insertion did not detectably impair entry into polysomes (**fig. S1C**).

We generated pegRNA libraries encoding diverse insertions to all three target sites and performed 10 serial rounds of transfection of PEmax-expressing K562 cells at four-day intervals, measuring barcoding efficiency at each round (**Fig. 1D**). Most editing occurred in the first few rounds, and bulk efficiencies reached 93%, 91% and 80% by day 40 with pegRNAs targeting 18S, ITS1 and 28S, respectively (**Fig. 1E**). To establish a one-to-one correspondence between barcodes and units, we derived 16 monoclonal lines per target site. Clonal barcoding rates averaged 94%, 92% and 83%, matching the pools from which they derived, and the best clone at each site reached 98%, 95% and 99% (**Fig. 1F**). These clones carried approximately 52, 99 and 135 distinct barcodes (**fig. S1D-E**). For genome assembly, we selected a 28S-barcoded clone in which 99% of reads overlapping the target site carried a barcode, as it combined the highest barcoding rate and the greatest barcode diversity at a ribosomally-retained, variation-tolerant target site.

### Barcode-guided assembly of 18 NOR arrays at single rDNA-unit resolution

To assemble NORs, we subjected gDNA from the selected 28S-barcoded K562 clone to nanopore sequencing, generating ∼900 Gb with a read N50 of 39.5 kb (Oxford Nanopore Technologies, ONT). We then applied a custom, barcode-guided assembly approach (**fig. S2A**; **Methods**). <u>First</u>, we extracted reads containing rDNA barcodes (n = 13,711; N50: 68.3 kb) and, after removing palindromic reads (n = 7), classified them into single-barcode (n = 11,571; N50: 45.5 kb), 2-barcode (n = 1,425; N50: 88.6 kb) and ≥3-barcode (n = 708; N50: 171.8 kb) read groups. The longest read was 1.38 Mb and spanned a full NOR, including 10 barcoded rDNA units (**fig. S2B**; **table S1**). <u>Second</u>, to identify reads spanning the ends of NOR arrays, we used exact 20-mer matches to an rDNA reference sequence (KY962518.1) to discriminate rDNA vs. non-rDNA regions within reads, and then selected reads with >25 kb of non-rDNA sequence at either or both flanks (n = 418 in the single-barcode group; n = 189 in the multi-barcode groups). Collapsing redundant flanking sequences using multi-barcode reads and incorporating support from single-barcode reads, we obtained 18 upstream (5’) and 18 downstream (3’) flanking sequences, each supported by ≥4 reads (**fig. S2C**; **table S2**). <u>Third</u>, we performed assembly from the barcode orders observed within ≥3-barcode reads, which yielded 18 high-confidence NOR scaffolds, each anchored by one upstream and one downstream flank. This number was confirmed by fluorescence *in situ* hybridization (FISH) on metaphase spreads of this clone (*18*), which resolved 18 rDNA arrays across 17 chromosomes (**fig. S3A-B**). <u>Finally</u>, we aligned each supporting read to its scaffold (median 20 reads supporting each barcode position across all scaffolds; range, 3-34; IQR, 16-24; **Fig. 2A-B**), split the alignments at barcodes, and called a consensus for each resulting interval (**fig. S2D**). Concatenating consensus sequences in the barcode-supplied order yielded our final NOR assemblies (**data S1**).

**Figure 2.**
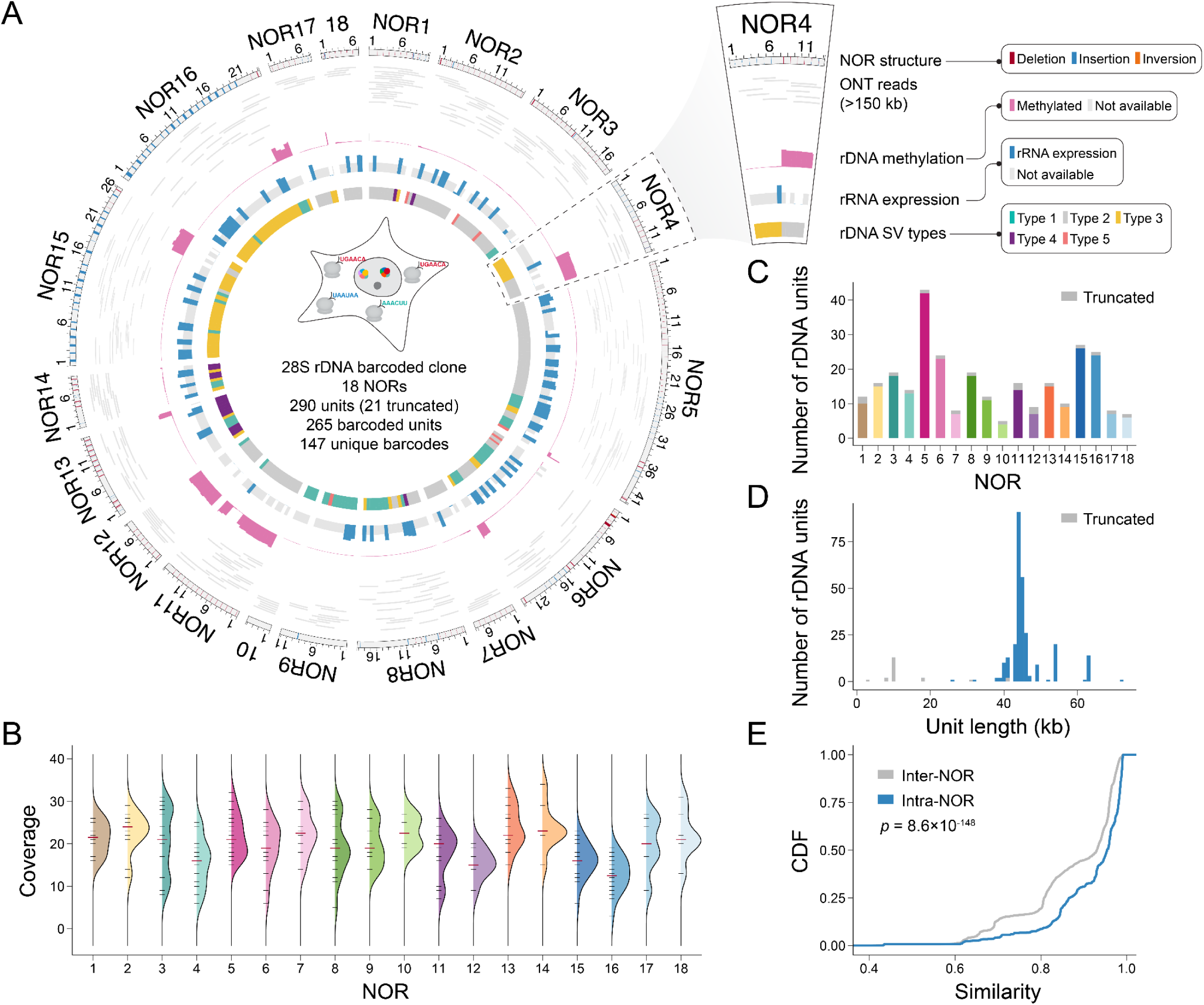
Barcode-guided assembly of 18 NOR arrays at a single rDNA-unit resolution. **(A)** Ring plot summarizing the 18 assembled NORs of the K562 genome. NORs are numbered by chromosomal location (see Fig. 5), so that related arrays appear adjacent. All rings are registered to the same rDNA units, delimited by black ticks in the outer ring. Outer ring, NOR architecture, with black ticks separating individual rDNA units and structural variants >500 bp colored by class (red, deletion; blue, insertion; orange, inversion). Second ring, ONT ultra-long reads (>150 kb) aligned to each assembly. Third ring, DNA methylation status across the 47S pre-rRNA region of each unit (see Fig. 6). Units for which methylation could not be assigned are grey. Fourth ring, transcriptional status of the rDNA units, based on barcode abundance in short-read sequencing of rRNA (see Fig. 6). rDNA units with non-unique barcodes and units lacking a barcode are grey. Innermost ring, structural variant type of each unit, as assigned by the region where the SV occurred (see Fig. 3). Truncated terminal units <35 kb are excluded and not shown in this ring. **(B)** Read coverage at barcode positions across the 18 NOR scaffolds. Short black ticks show coverage at individual barcode positions; the long red tick marks the median across each NOR’s barcodes. Curves to the right of each set of ticks are kernel density estimates of the coverage distribution within that NOR. **(C)** Number of rDNA units per NOR (n = 290 total; range 5-43). Truncated terminal units are shown in grey. **(D)** Length distribution of all 290 rDNA units; truncated units shown in grey. **(E)** Cumulative distribution functions (CDFs) of pairwise sequence similarity for pairs of full rDNA units from the same NOR (intra-NOR; n = 2,641 pairs) or different NORs (inter-NOR; n = 33,944 pairs). *p*-value from Wilcoxon rank-sum test.

Altogether, the 18 NOR scaffolds contained 290 full or partial rDNA units, of which 265 (91.4%) were tagged by one of 147 distinct barcodes (**Fig. 2C**). Of the 25 untagged units, 4 had an intact pegRNA target site, while 21 lacked it due to truncation or internal deletion. Thus, 265 of 269 (98.5%) of taggable rDNA units were indeed tagged. Of the 147 barcodes, 104 occurred once, 28 twice, and 14 three to seven times (**fig. S2E**). One (TGAACA) was a conspicuous outlier, with 44 appearances spanning all 18 NORs, a frequency unexplained by its representation in the plasmid library (**fig. S1E**). Inspection of ONT reads identified a stably integrated plasmid encoding this barcode, such that the cells’ cumulative exposure to this pegRNA far exceeded that of any other. Among the other 42 recurring barcodes, we observed 1 “run of three” consecutive rDNA units sharing a barcode and 3 “runs of two”, against permutation-based expectations of 0.008 ± 0.087 and 0.756 ± 0.847 (**fig. S2F-G**). Adjacency is the signature expected of unequal crossover or gene conversion, and we speculate that the handful of duplicate instances that appear adjacently arose from ongoing homogenization during the construction and passaging of this clone, rather than from independent prime editing events.

To define rDNA unit boundaries consistently, we set the unit start ∼3.7 kb upstream of the 5′ ETS, where 20-mers transition from unique to repetitive in 15 of the 18 NORs, and re-indexed the reference sequence and assemblies accordingly. Oriented 5′ to 3′, the remaining three NORs (NORs 1, 11, 12) begin with truncated units, at two distinct breakpoints, while all 18 NORs end with truncated units. Fifteen of these 3′ breakpoints are close to those reported in other T2T genomes and span the 18S to ITS1 regions (*10*, *11*). The other three (NORs 2, 7, 18) end at two further breakpoints: one unique to NOR2, and one shared by NORs 7 and 18 (**table S3**). Below, we assign each NOR to its chromosome, which shows that these atypical ends coincide with chromosomes rearranged during the history of K562. Unit lengths spanned 2.8 to 72 kb but were tightly concentrated (median 44.3 kb; IQR 44.0-45.8 kb; **Fig. 2D**), with the tails contributed by truncated terminal units and by units carrying large IGS rearrangements. Excluding 19 truncated terminal units (those <35 kb; **table S3**), pairwise comparison of the remaining 271 revealed extensive heterogeneity, with intra-NOR similarity exceeding inter-NOR similarity (median 95.4% vs. 93.0%; *p* = 8.6 × 10^-148^, Wilcoxon rank-sum test), consistent with partial homogenization within arrays (**Fig. 2E**).

### Analysis of rDNA sequence heterogeneity

We next characterized sequence heterogeneity among these 271 rDNA units (*i.e.* continuing to exclude truncated terminal units; **table S3**). The rDNA reference sequence (KY962518.1; 44,838 nt) comprises a 13-kb Pol I transcription cassette encoding the 47S pre-rRNA, followed by an IGS containing three R repeats and two Butterfly/Long repeats (*4*). Aligning each unit to this reference revealed pervasive structural variants (SVs), predominantly deletions and amplifications of the R and Butterfly/Long repeats, and occasionally rearrangements impacting the 47S pre-rRNA itself (**data S2**).

To group units systematically by such structural variation, we performed principal component analysis (PCA) on their 6-mer content (**fig. S4A**), and then partitioned the units by *k*-means clustering (*k* = 6; **Fig. 3A**). PC1 accounted for 58% of the variance, and the 6-mers most strongly correlated with PC1 were enriched in the Butterfly/Long and R repeats of the IGS (**fig. S4B**). Consistent with this, units assigned to clusters 3, 1, 2 and 4 are ordered along PC1 by a progressive increase in Butterfly/Long repeat number (**Fig. 3A**; **fig. S4C**). As each Butterfly/Long repeat is ∼4.5 kb, this variation accounts for the non-canonical modes of rDNA unit lengths at 41, 49, 54, 63 and 72 kb (**Fig. 2D**). Units of clusters 5 and 6 carry SVs affecting the 47S pre-rRNA itself, and separate from the rest along PC2 (**fig. S4C**).

**Figure 3.**
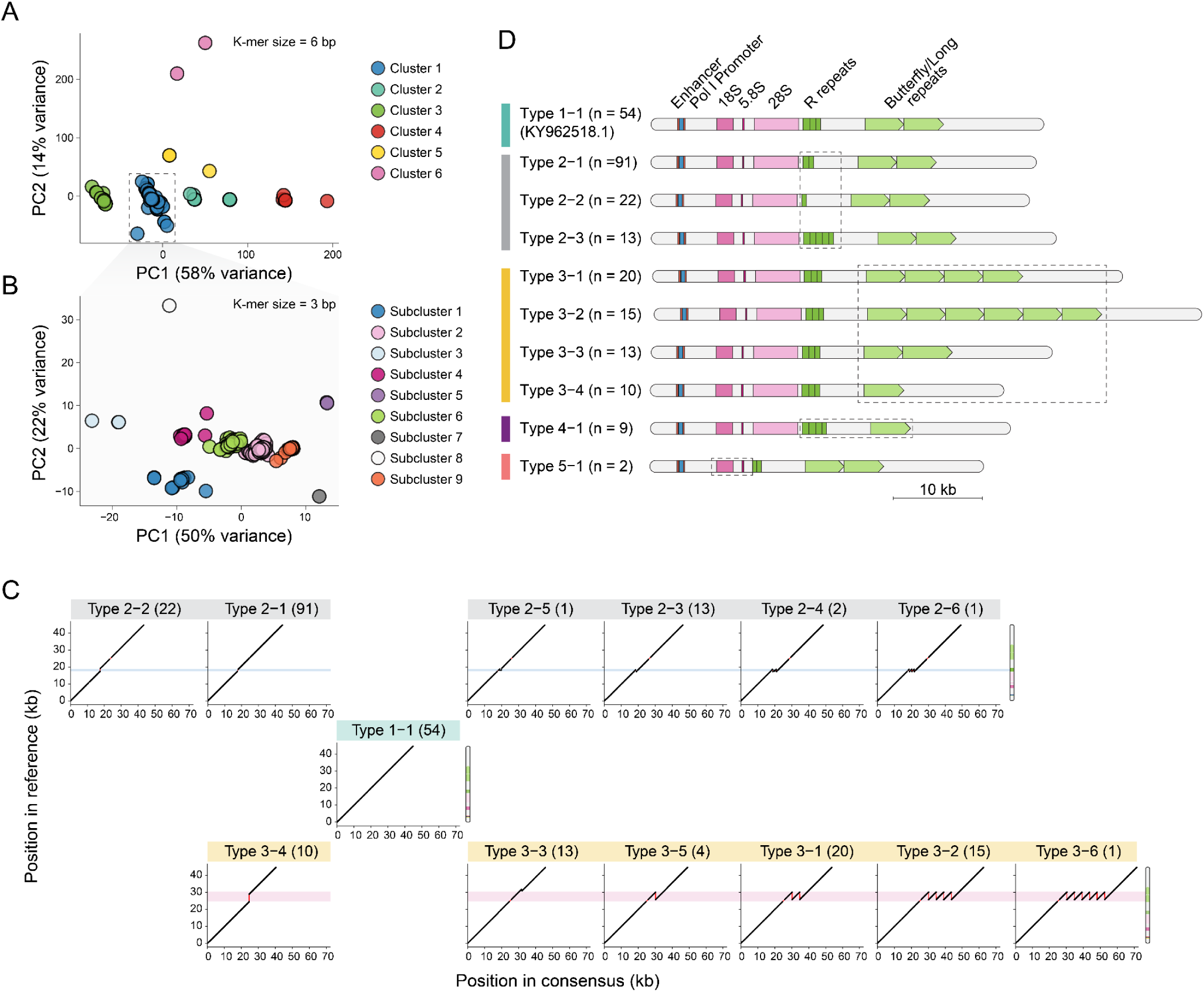
Analysis of rDNA sequence heterogeneity. **(A)** PCA of the 271 rDNA units based on 6-mer content, showing PC1 and PC2. Clusters were defined by *k*-means clustering (*k* = 6). **(B)** Units in cluster 1 (n = 201) were subjected to a second round of PCA based on 3-mer content, showing PC1 and PC2. Subclusters were defined by *k*-means clustering (*k* = 9). **(C)** The clusters in **A** and **B** were manually curated into five primary types, defined by whether SVs affect the R repeats, Butterfly/Long repeats or 47S pre-rRNA (**Methods**; **fig. S4E**). Primary types 2-5 were further curated into 6 subtypes, such that together with Type 1-1, there are a total of 25 labels (**table S4**). Shown is the sequence alignment between the consensus sequence of each group in the first three primary types (x-axis) and the reference sequence KY962518.1 (y-axis). Within each type, subtypes are numbered in descending order of unit count. Subpanels are manually arranged so that subtypes sharing SV-affected regions can be highlighted by horizontal colored strips, which project onto a schematic of the reference allele at right. **(D)** Simplified schematics of the reference rDNA allele (KY962518.1), which matches Type 1-1, together with one or several representatives of each of the other primary types. Dashed boxes highlight regions altered by SVs relative to the reference.

Because most rDNA units fall within cluster 1 (n = 201), we repeated k-mer-based PCA on these units only, now with 3-mers rather than 6-mers counts, followed by *k*-means clustering (*k* = 9; **Fig. 3B**). Units assigned to subclusters 3, 4, 6, 2 and 9 fall along the new PC1 and track a progressive decrease in the length of the R repeat region (*55*). Units in subcluster 1 bear a small duplication within one Butterfly/Long repeat, while units of subcluster 8 carry rearrangements in the 47S pre-rRNA region (**fig. S4D**).

Together, these analyses show that rDNA units in the K562 genome can be grouped based on their SV in three regions: the R repeats, the Butterfly/Long repeats, and the 47S pre-rRNA. Starting from the 14 groups defined by the two rounds of k-mer-content-based clustering, we manually curated 5 primary types of rDNA units, 4 of which could be broken down into 6 subtypes (25 labels altogether; **Fig. 3C**; **fig. S4E**; **table S4**). Briefly, these are: (i) <u>Type 1</u>: 54 of 271 (20%) rDNA units that bear two Butterfly/Long and three R repeats, resembling the reference allele; (ii) <u>Type 2</u>: 130 (48%) that bear varying degrees of deletions or amplification of their R repeats. A subtype of these, Type 2-1, characterized by a deletion of a single R repeat relative to the KY962518.1 reference, is the most abundant class across all labels (n = 91); (iii) <u>Type 3</u>: 63 units (23%) bear varying numbers of Butterfly/Long repeats (median 4, IQR 2-6, range 1-8); (iv) <u>Type 4</u>: 17 units (6%) bear duplications and/or deletions involving both the Butterfly/Long and R repeats; and (v) <u>Type 5</u>: 7 units (3%) bear rearrangements impacting the 47S pre-rRNA region (**Fig. 3C-D**).

To assess which of these SV “types” are also present in germline human genomes, we projected rDNA units from T2T reference genomes onto the same PCA space (*10*, *11*) (**fig. S5A-B**). Across all three genomes, Type 1 or Type 2-1 were the most common configurations, accounting for 54% of units in K562 (145 of 271), 45% of unique sequences in CHM13 (15 of 33), and 68% in HG002 (27 of 40) (**Methods**; **fig. S5C-D**). Of the 31 remaining other T2T configurations, 29 were distributed across nine other K562 types (2-2, 2-3, 2-4, 2-5, 3-3, 3-5, 4-1, 4-4 and 4-6), while one combined features of Types 2-2 and 3-3, and one combined features of Type 2-2 and 3-5 (**fig. S5D**; **table S4**). The types defined here therefore account for essentially all rDNA configurations observed in CHM13 and HG002, with the caveat that only a subset of their full diversity may be represented in the T2T assemblies. Conversely, the rarer types we observe in K562 appear to be additions to, rather than substitutes for, the configurations found in germline genomes, and plausibly arose during cancer genome evolution or the passaging of this cell line.

### *Cis-a*rchitecture and chromosomal locations of the 18 NORs of the K562 genome

To visualize the distribution of rDNA unit types along each NOR, we encoded the five primary types by color and their subtypes by symbol (**Fig. 4A**). Striking patterns were immediately evident on visual inspection. Within NORs, units of a given type are not distributed randomly but rather occur in contiguous runs, as expected if unequal crossover and gene conversion homogenize units locally. To compare arrays quantitatively, we computed a pairwise similarity score between every pair of NORs, based on the mean sequence similarity of reciprocal best hits of their constituent rDNA units (**Methods**). Agglomerative hierarchical clustering on this matrix identified seven groups of highly related arrays: NORs 1, 17, 7, 18; NORs 2, 3; NORs 5, 6; NORs 9, 10; NORs 11, 12; NORs 13, 14; and NORs 15, 16 (**Fig. 4B**). Two arrays, NOR4 and NOR8, were relatively distant from all others. An independent, k-mer-based metric computed at the whole-array level resulted mostly in similar groupings (**fig. S6A**).

**Figure 4.**
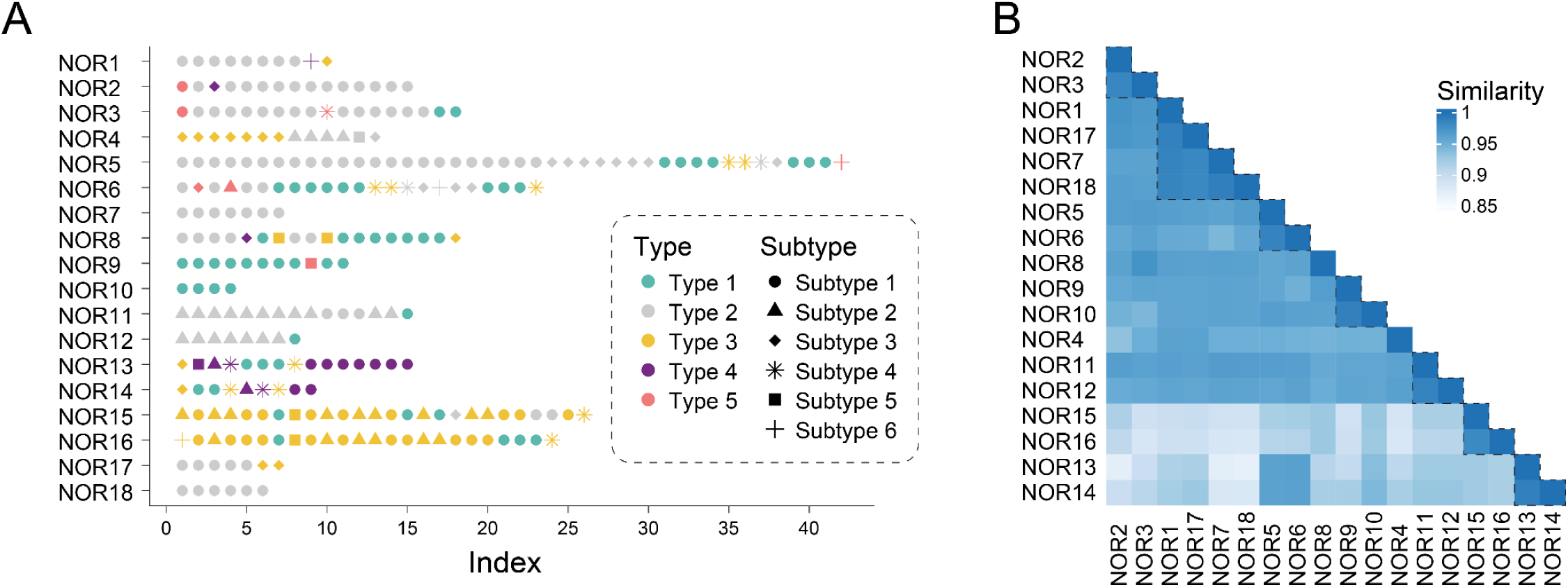
*Cis-a*rchitecture and pairwise similarities across the 18 NORs of the K562 genome. **(A)** Structural variant type of each rDNA unit (n = 271) and its position within each of the 18 NORs. The five primary types are color-coded and their subtypes indicated by distinct shapes. Truncated terminal units less than 35 kb are not shown. **(B)** Pairwise similarity between NORs. For each unit of one NOR, the highest sequence similarity to any unit in the other NOR was identified, and these best-hit similarities were averaged across all units. This procedure was performed reciprocally, and the two means then averaged to give the plotted pairwise similarity. NORs were clustered on the similarity matrix by agglomerative hierarchical clustering (Ward’s minimum-variance method, Euclidean distance) and displayed as a heatmap. Diagonal values were set to 1. Brackets mark the 7 groups of putatively related NORs discussed in the text.

A diploid, germline human genome contains 10 NORs on the short arms of chrs. 13, 14, 15, 21 and 22. In contrast, we identified and assembled 18 NORs in the pseudotriploid genome of K562, a BCR-ABL-driven chronic myelogenous leukemia line carrying the Philadelphia chromosome, t(9;22)(q34;q11) (*52*). To determine the chromosomal location of each NOR, we performed whole genome assembly of K562 with *Hifiasm* (*56*, *57*) (**Methods**; **table S1**). We then located each NOR by aligning its flanking sequences to this assembly, further guided by FISH (**fig. S3B**). The result was a one-to-one mapping of NORs to host chromosomes in which rDNA probe intensity correlated with NOR size (**Fig. 5**; **fig. S3C**).

**Figure 5.**
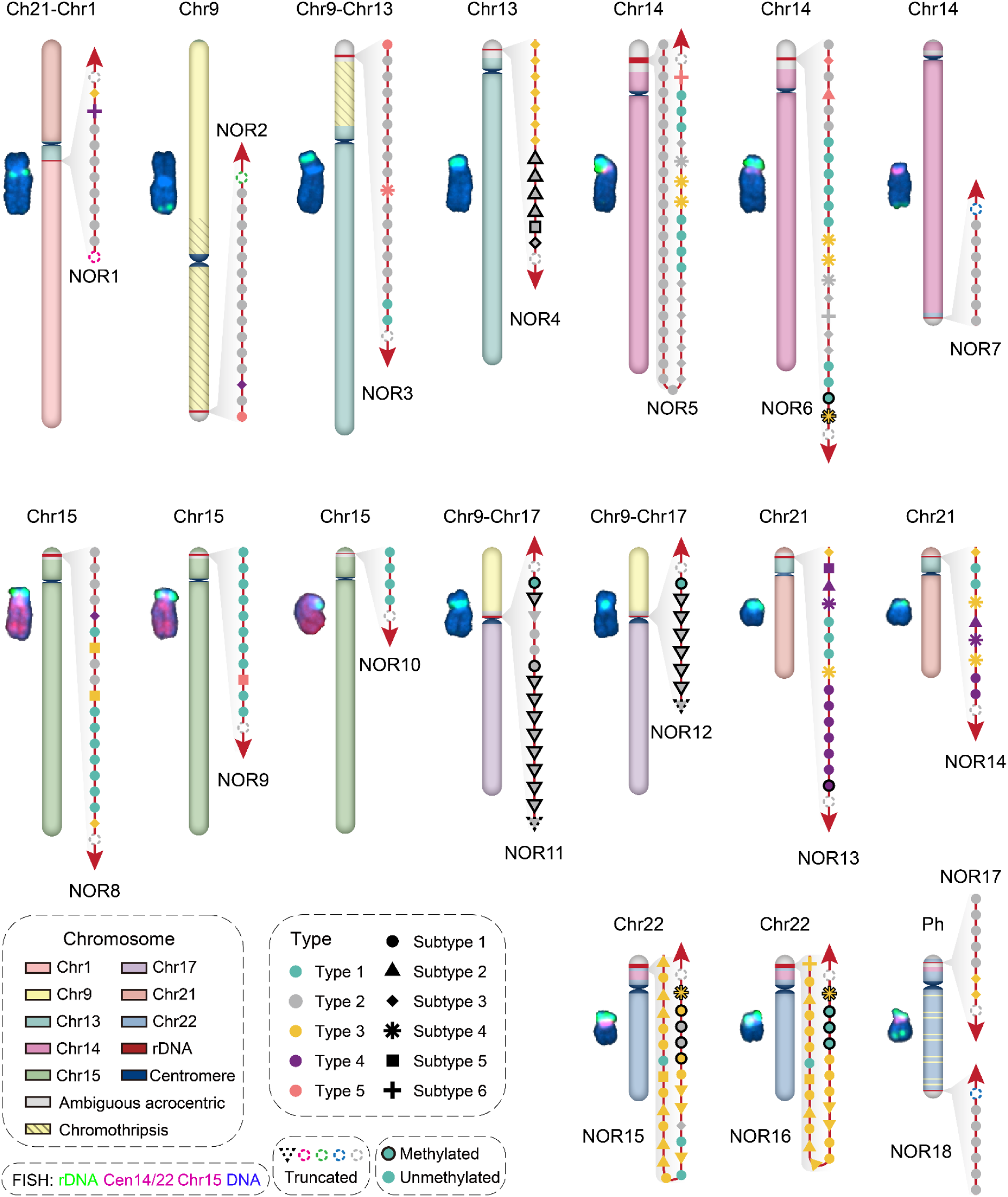
*Cis-a*rchitecture and chromosomal locations of the 18 NORs of the K562 genome. Schematic structures of the 17 NOR-bearing chromosomes of the K562 genome, with matched FISH images shown alongside. Scaffolds are annotated by sequence alignment to the CHM13 genome. Alignments within the acrocentric short arms that were low-confidence or ambiguous are shown in grey (**Methods**). In the FISH images shown to the left of each chromosomal schematic, rDNA is labeled in green; the centromeres of chrs. 14 and 22 and the whole of chr15 are labeled in red. For each chromosome, the organization of rDNA units within its NOR is shown at its assigned position in the assembly. The 5’→ 3’ orientation of each NOR (as per our nomenclature) is indicated by the red arrowheads; note that with the exception of NOR11 and NOR12, the 5’ end of each NOR is positioned closer to the telomere, and the 3’ end closer to the centromere. The five primary unit types are color-coded and their subtypes indicated by distinct shapes, as in Fig. 4A. Truncated terminal units are indicated by dashed outlines, with dash color denoting the truncation breakpoint class (**table S3**). Units with high 47S methylation are outlined in black (see Fig. 6).

Ten NORs mapped to the short arm of karyotypically intact, acrocentric chromosomes, including one singleton (NOR4 on chr13), three pairs (NORs 5, 6 on chr14; NORs 13, 14 on chr21; NORs 15, 16 on chr22) and one triplet (NORs 8, 9, 10 on chr15). The 5’ ends of each of these NOR (as per our orientation nomenclature) were consistently positioned closer to the telomere and the 3’ ends closer to the centromere (**Fig. 5**). All three pairs appear to correspond to chromosomal duplications followed by independent divergence of the two copies, as evidenced by high pairwise similarity of the NORs and patterns of loss-of-heterozygosity (LOH) (**Fig. 4B**; **fig. S6A,C**). Within the chr15 triplet, NOR9 and NOR10 also appear to derive from chromosomal duplication, while NOR8 appears to correspond to the germline homolog, as evidenced by its divergence from NORs 9 and 10 and allelic heterozygosity ratios (**Fig. 4B**; **fig. S6A,C**).

The remaining eight NORs reside on rearranged chromosomes, and their positions trace the history of this cancer genome (**Fig. 5**). In K562, the Philadelphia chromosome carries a focal amplification of the BCR-ABL fusion and is larger than the classical form (*51*). It harbors two NORs, one at the chr22 acrocentric p arm (NOR17) and one near the end of its q arm (NOR18). This founding event appears to have destabilized chr9, which participates in additional rearrangements, including three involving four NORs: (i) a duplicated chr9-chr17 fusion [der(17)t(9;17)(9pter→9p21::17p11→17qter)] (*51*) carries an array near its centromere, accounting for the highly related NOR11 and NOR12; (ii) a chromosome dominated by rearranged chr9 fragments, consistent with chromothripsis, contains NOR2; (iii) a chr9-chr13 fusion [der(13)t(9;13)(9q12::13p11→13qter)] (*51*) carries an enlarged chr13 “p arm” bearing NOR3 with high similarity to NOR2, suggesting a shared origin subsequent to the putative chromothripsis event (**Fig. 4B**; **fig. S6A**).

Two further translocations impacted rDNA without duplicating it. NOR1 remains at its original chr21 position but is truncated at its 5′ end by a translocation with chr1 [der(21)t(1;21)(1qter→1q23::21p11→21qter)] (*51*). NOR7 sits at the end of the chr14 q arm rather than on an acrocentric short arm. Its closest relative by sequence is NOR18, on the Philadelphia chromosome (**Fig. 4B**; **fig. S6A**), with which it also shares a terminal truncation breakpoint (**table S3**), suggesting that the two share a common origin rather than NOR7 representing a relocated chr14 p-arm NOR.

Together, these assignments show that the expansion from 10 NORs (germline) to 18 NORs (K562) arose by two distinct routes. Whole-chromosome duplications produced five pairs of arrays -- NORs 5, 6; NORs 9, 10; NORs 11, 12; NORs 13, 14; and NORs 15, 16 -- that remain highly similar but have diverged measurably since duplication, offering a natural experiment in the rate and mode of rDNA array evolution. Separately, translocation and chromothripsis moved rDNA arrays to non-acrocentric positions, including the q arms of the Philadelphia chromosome and of chr14, indicating that NORs are not merely passive casualties of cancer genome instability, but are relocated and truncated by it, and may even contribute to it as substrates for rearrangement. That these events are legible at all reflects the sequence heterogeneity retained within and between arrays. Despite the homogenizing effect of recombination, individual NORs remain distinguishable, and their relationships recoverable.

### Individual rDNA units adopt stable, binary methylation and expression states

Beyond enabling NOR assembly, the barcodes introduced by prime editing serve as stable labels of individual rDNA units, read out by any sequencing modality applied to these cells. We sought to leverage this feature to explore patterns of DNA methylation and rRNA expression, as well as the extent to which these are correlated, across individual rDNA units and assembled NORs.

rDNA methylation is inversely correlated with rRNA output and has been proposed as its key determinant (*2*). Nanopore sequencing of single DNA molecules has further revealed an “all-or-none” methylation pattern for individual rDNA units (*3*). However, absent a unit-resolved map, it has not been possible to ask how methylation is distributed across NORs, nor whether methylation states of individual rDNA units are stable across cell divisions. To explore the stability of rDNA methylation and its relationship to transcription at single unit resolution, we extracted CpG methylation (5mC) calls from ONT reads and computed methylation profiles for each of the 271 rDNA units (**Methods**). Units fell into two discrete states (**Fig. 6A**), with 85% (n = 231) lowly methylated across the 47S pre-rRNA region but highly methylated in the IGS (“low-methylation”), while 14% (n = 38) showed the reverse configuration, highly methylated across the 47S but lowly methylated in the IGS (“high-methylation”). The remaining two units, which bear a near-complete deletion of the 47S pre-rRNA region (Types 5-2 and 5-3), have methylated IGS regions (**Fig. 6B**; **fig. S7A**; **data S3**).

**Figure 6.**
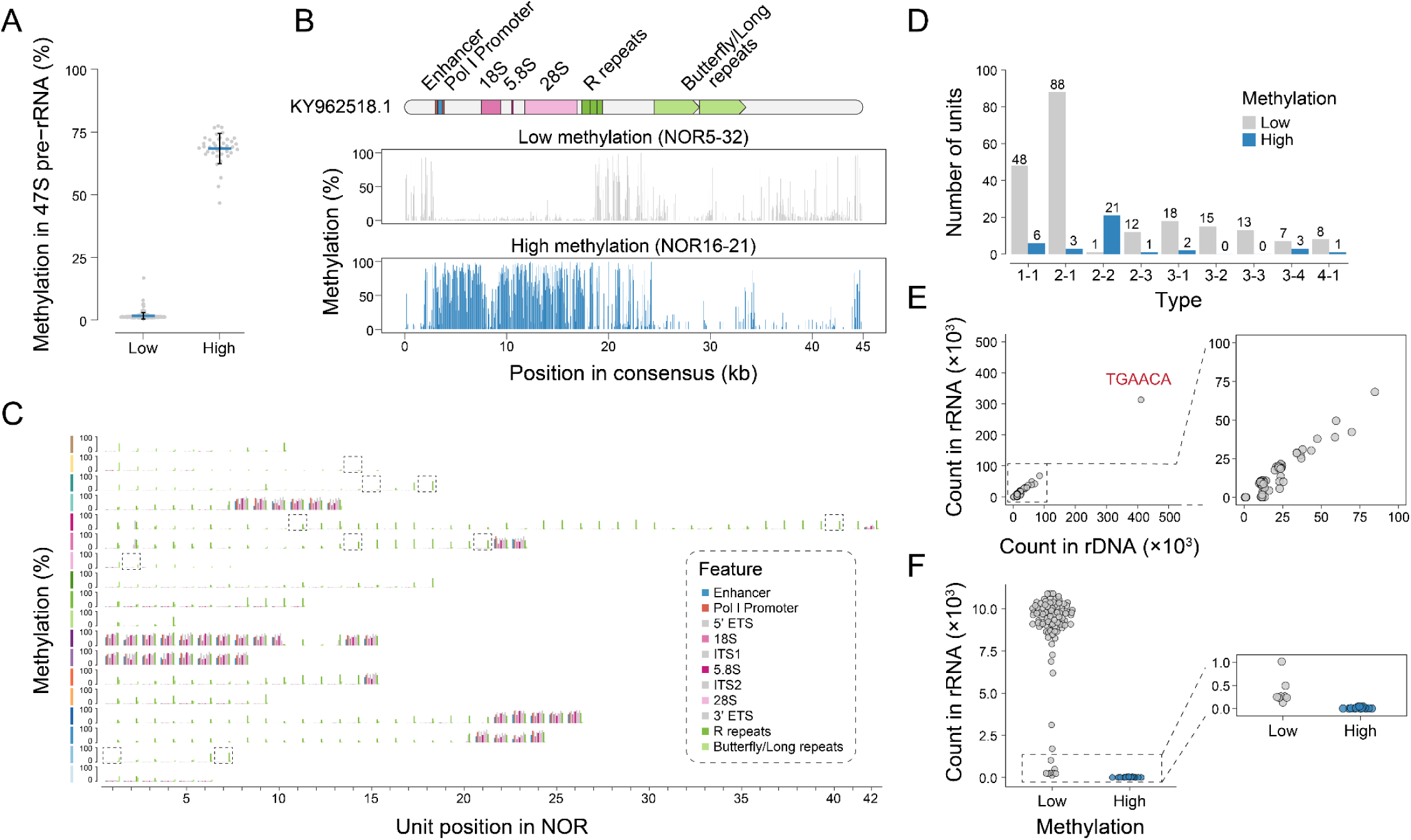
Individual rDNA units adopt stable, binary methylation and expression states. **(A)** Methylation (5mC) levels, averaged across the 47S pre-rRNA and the upstream regulatory region for the 269 rDNA units (*i.e.* excluding the two units with a nearly complete deletion of this region), are bimodally distributed: 231 units with low 47S methylation (median 1.4%; range 0.5-17%; IQR 1.2-1.8%) and 38 with high 47S methylation (median 69%; range 47-77%; IQR 67-72%). **(B)** Representative methylation profiles across the full rDNA unit, for one exhibiting low 47S methylation (top) and one high 47S methylation (bottom), both Type 1-1. **(C)** Methylation levels, averaged within each major region of each rDNA unit, plotted by NOR (rows) and unit position (columns). Dashed boxes mark lowly expressed but low-methylation units with unique barcodes (count in rRNA < 5,000 in **F**). **(D)** Bar plot showing 47S pre-rRNA methylation status for the most common rDNA subtypes. **(E)** Correlation of rDNA barcode abundance in short-read sequencing between gDNA (x-axis) and total RNA samples (y-axis). The point corresponding to barcode “TGAACA” is annotated. Spearman’s *ρ* = 0.73. **(F)** RNA abundance of low 47S methylation units (left) versus high 47S methylation units (right), for the 104 barcodes present in a single rDNA unit.

Because each profile averages dozens of ONT reads originating from different cells, the binary distribution implies that methylation state of individual rDNA units is stably maintained across cell divisions. Mapping these states back onto the NOR assemblies, the high-methylation units were non-randomly distributed, concentrated toward the ends of arrays (**Fig. 2A**; **Fig. 6C**). They were also unevenly distributed across SV types: 21 of 22 Type 2-2 units were high-methylation, which may reflect their position near the centromere of the chr9-chr17 fusion (NORs 11 and 12; **Fig. 5**; **Fig. 6D**).

To relate methylation to transcription, we measured rDNA barcode abundance in both gDNA and total RNA from the same cells, taking the latter as a proxy for rRNA output. Barcode abundances were well correlated between the two (*ρ* = 0.73; **Fig. 6E**). We then restricted the analysis to the 104 unique barcodes, whose assignments to specific rDNA units are unambiguous. Expression remained binary (**Fig. 6F**). Furthermore, expressed units were all low-methylation and transcribed at comparable levels, while high-methylation units were barely detectable in rRNA, confirming the link between rDNA methylation and silencing (**Fig. 6F**). Surprisingly, 10 rDNA units were lowly expressed despite being in a low-methylation state (**Fig. 6C,F**). There was no detectable difference in methylation between expressed and unexpressed low-methylation units (**fig. S7B**), indicating that methylation of the 47S region is sufficient but not necessary for rDNA silencing. These ten units are distributed across six NORs at both terminal and internal positions (**Fig. 6C**; **fig. S7C**) and span four abundant SV types (1–1, 2–1, 3–3, 3–4). What factors underlie their lack of expression remains unclear.

## DISCUSSION

NORs, arguably the darkest of the genome’s “dark matter”, remain incompletely resolved in every human genome assembly to date, hindering our understanding of these fundamentally important regions. Here we present a fully resolved genetic and epigenetic map of every NOR in the pseudotriploid genome of a widely used human cancer cell line, K562, at single-unit resolution. To achieve this, we wrote distinct DNA barcodes into the highly homologous rDNA units in a saturating manner that allowed individual units to be distinguished in both long and short reads, and at both the genomic and transcriptomic levels. Our assemblies reveal the cis- and trans-architecture of sequence variation across rDNA units and NORs, along with patterns of NOR organization that have been actively shaped by cancer genome evolution. Finally, we link the binary pattern of transcriptional activity of rDNA units to their methylation status, confirming the expected anticorrelation while also discovering that some rDNA units remain lowly expressed despite low methylation in the 47S region.

Complete assembly of K562’s NORs enabled a comprehensive assessment of rDNA sequence heterogeneity. Principal component analysis of 6-mer and 3-mer frequencies revealed distinct, visually separable clusters of units, most corresponding to deletions or amplifications of the Butterfly/Long and R repeats. Of 271 full rDNA units, 119 (44%) carry three R repeats and 189 (70%) carry two normal Butterfly/Long repeats, matching the reference allele (KY962518.1). Because Butterfly/Long copy number varies on a three-R background (Type 1 → Type 3), while R copy number varies on a two Butterfly/Long repeat-background (Type 1 → Type 2), we suspect that the three-R, two-Butterfly/Long configuration may be the ancestral state from which other types derive (**Fig. 3D**).

The K562 cell line was established in 1975, began to acquire a partially triploid karyotype by 1983, and eventually evolved into a pseudotriploid state (*51*, *52*, *58*). That we resolve 18 NORs, rather than the 10 expected for a germline diploid genome, follows from this history. Ten NORs reside on the short arms of intact acrocentric chromosomes: NOR4 (chr13), NORs 5, 6 (chr14), NORs 8, 9, 10 (chr15), NORs 13, 14 (chr21), and NORs 15, 16 (chr22). NORs 5 and 6, 9 and 10, 13 and 14, and 15 and 16 appear to derive from chromosomal duplication followed by independent divergence, while NOR8 appears to be the germline homolog of NORs 9 and 10. The remaining eight reside on rearranged chromosomes: NORs 17 and 18 on the Philadelphia chromosome; NORs 11 and 12, a fifth duplicated pair, on a duplicated chr9-chr17 fusion; the highly similar NOR3 on a chr9-chr13 fusion and NOR2 on a chr9-derived chromosome whose structure is suggestive of chromothripsis; NOR1 on a chr1-chr21 translocation; and NOR7 at the end of a chr14 q arm. Thus, the expansion from 10 to 18 NORs arose by whole-chromosome duplication, which produced five pairs of arrays, and by translocation and/or chromothripsis, which relocated and truncated arrays. The persistence of the five duplicated pairs (NORs 5/6, 9/10, 11/12, 13/14 and 15/16) as distinct entities offers a natural experiment in how rDNA arrays diverge over time, although the timing of the duplications is unknown. NORs 2 and 3, which appear to share a common chr9-associated origin, plausibly represent another such pair.

Although rDNA methylation has long been associated with rRNA silencing, its stability over time and its effects at the level of individual units have remained elusive. The K562 cells characterized here were derived from a single clone, which allowed us to assess the stability and distribution of this epigenetic mark in a uniform genetic background. After collapsing ONT reads for the same unit across progeny of the founder clone, we observe a clear binary distribution of methylation profiles. This suggests the epigenetic state remained stable across at least roughly 30 doublings, estimated from the number of passages and cells harvested for ONT sequencing. Among structural variants, Type 2-2 was enriched for high-methylation units. A recent study reported a correlation between low R repeat number with high rDNA methylation on the chr22 of CHM13 (*18*). However, this enrichment could reflect expansion of a single ancestral high-methylation unit, which makes it difficult to infer a causal relationship between rDNA sequence variation and methylation status. The expression of rRNA from individual rDNA units followed an “all-or-none” pattern (*59*), with methylation of the 47S pre-rRNA region tightly associated with silencing. Interestingly, although high-methylation units were barely detectable in rRNA, ten low-methylation units were also lowly expressed, suggesting that methylation is sufficient but not necessary for silencing.

Saturation barcoding was critical to the success of our NOR assemblies. By leveraging the unique chromatin and transcriptional status of rDNA, together with successive rounds of transfection, we achieved a 99% barcoding rate, which, to our knowledge, has not previously been reported for prime editing. Of course, this raises the concern that the editing itself may cause genome instability, including rDNA loss. Although we cannot rule out such alterations, the karyotype of the NOR-bearing chromosomes largely recapitulates the K562 karyotype reported in 2001 (*51*). We also note that we benefited here from K562’s high prime editing efficiency, and caution this approach may be more challenging in cell lines less amenable to prime editing.

Finally, the ease with which NORs were prime edited is itself notable. Because NORs comprise hundreds of redundant, actively transcribed copies, they may be a particularly permissive site for certain kinds of genome editing. In the clone used here, 265 units, or 99% of all units bearing the targeted spacer, were successfully edited. The counterargument is that rDNA arrays are recombinationally dynamic, and unequal crossover or gene conversion could delete or homogenize edited sequences over time, as the adjacent duplicate barcodes we observed suggest. NORs may therefore be most useful where multicopy genome editing is desired and loss or gain of some copies is tolerable, rather than as a stable single-copy safe harbor.

In summary, we demonstrate a means of resolving NORs, the “final mile” in human genome assemblies, and shed light on their complex genetic and regulatory landscape. Although this is a single snapshot of a single cancer genome, our results, together with other recent work (*18*), suggest that NORs are remarkably dynamic. They diverge substantially between germline homologs, are reshaped by structural rearrangements in cancer, and may contribute to those rearrangements. Looking forward, barcoded and assembled NORs may enable the deeper investigation of rDNA evolution, epigenetic regulation, and the consequences of unit-level variation for nucleolar organization and ribosome biogenesis, and may even allow specific ribosome subtypes to be linked to the mRNAs they translate. More broadly, prime-editing-based barcoding offers a general strategy for resolving complex repetitive regions, and could be extended to introduce defined rDNA sequence and structural variants to test their effects on molecular and organismal phenotypes.

## Acknowledgments

We thank A. Phillippy (JHU), M. Vollger (UW), D. Antipov (JHU), all members of the Shendure Lab at the University of Washington, and all members of the Li Lab at Westlake University for discussions and feedback. We also thank R. Waterston and C. Queitsch for their detailed feedback on the manuscript. This work was supported by the Paul G. Allen Frontiers Group (Allen Discovery Center for Cell Lineage Tracing to J.S.), Seattle Hub for Synthetic Biology, a collaboration between the Allen Institute, Chan Zuckerberg Initiative and University of Washington (award number CZIF2023-008738 to J.S.), the Brotman Baty Institute for Precision Medicine (J.S.), Westlake University, and Zhejiang Provincial Natural Science Foundation (award number LQK26C060001 to X.L.). J.S. is an investigator of the Howard Hughes Medical Institute.

## Author contributions

Conceptualization X.L., J.W., and J.S.; methodology: X.L., J.W., S.D.Z., and J.S.; investigation: X.L., S.D.Z., and J.W.; formal analysis and software: S.D.Z. and X.L.; writing: X.L., J.S., and S.D.Z.; funding acquisition: J.S. and X.L.; resources: T.P., S.Z.Z., X.R.L., R.M.D., and J.L.G; supervision: J.S. and X.L.

## Disclosures

We disclose that language editing, proofreading and coding were supported by AI-based tools; these were not used for conceptual development or primary manuscript writing. The authors take full responsibility for the contents of this manuscript.

## Competing interests

J.S. is on the scientific advisory board, a consultant, and/or a co-founder of Guardant Health, Phase Genomics, Pacific Biosciences, Cellular Intelligence and 10x Genomics. All other authors declare no competing interests.

## Data and materials availability

The Oxford Nanopore Technology Ultra-long DNA sequencing results are available at https://www.ncbi.nlm.nih.gov/bioproject/PRJNA1371648. All other sequencing results are archived at https://www.ncbi.nlm.nih.gov/geo/ with Accession ID GSE343139. An integrated genome browser view of the NOR assemblies, including sequences, annotations, unit types, barcodes, expression, methylation, and SVs, is available at https://shudizhao.github.io/k562_NOR_browser. The code for NOR sequence assembly, methylation analysis, and structural variant (SV) detection is available at https://github.com/ShudiZHAO/k562_NOR.

## METHODS

### Cell culture conditions

K562 PEmax(+) cells (*49*) were maintained in RPMI 1640 medium (Gibco) supplied with 10% FBS (Hyclone) and penicillin/streptomycin (Gibco, 100 U/ mL). HEK293T PEmax(+) cells were maintained in DMEM medium (Gibco) supplied with 10% FBS and penicillin/streptomycin. All cells were kept in a humidified incubator at 37 °C, 5% CO_2_.

### pegRNA design

pegRNAs targeting rDNA were designed against an rDNA unit from chromosome 22 (*5*). gRNA sequences were selected using Flashfry based on the Doench2014OnTarget score (*60*). Sequences of the target loci of the top 100 gRNAs were extracted. To find the optimal pegRNA, edited sequences were designed as target sequences carrying a 3-nt insertion at the Cas9 cut site. Thus, for each target site, 64 pairs of wild-type and edited sequences were input into DeepPrime (*61*) to identify pegRNA with the highest predicted score. 11 pegRNAs for 11 distinct target sites were chosen, each making a specific 3-bp insertion to its target (**table S5**).

### Molecular cloning

#### 3-nt insertion epegRNA vectors

An empty vector containing a U6 promoter, SpCas9 gRNA scaffold, and evopreQ1 was digested with Esp3I and EcoRI-HF (NEB). PegRNAs were created by amplifying the gRNA scaffold with primers attaching the spacer and the 3’ extension sequence (p1-p22). The PCR product was purified and cloned into the backbone with NEB HiFi assembly.

#### rDNA barcoding epegRNA libraries

For the 18S and ITS1 rDNA barcoding libraries, the 3-nt insertion pegRNA vectors were amplified with phosphorylated primers p23/p24 or p23/p25, respectively. The PCR product was purified with AMPure XP beads (Beckman) and ligated with T4 ligase. The ligation product was concentrated with AMPure XP beads and electroporated into NEB 10-beta electrocompetent *E. coli*.

For the 28S rDNA barcoding library, the gRNA scaffold was amplified by p26 and p27 and purified. The PCR product was cloned into the Esp3I and EcoRI-HF digested backbone by NEB HiFi assembly. The assembly product was concentrated with AMPure XP beads and electroporated into NEB 10-beta electrocompetent *E. coli*.

A full list of oligos used can be found in **table S6**.

### General transfection strategies

Transfections of K562 cells were performed using a Lonza Bioscience 4D-Nucleofector system and the SF Cell Line 96-well Nucleofector® Kit (Lonza). For each well (20 μL), 2 × 10^5^ ∼ 4 × 10^5^ cells were transfected with up to 1.2 μg of DNA. The program code was FF-120.

Transfection of HEK293T cells was performed with Lipofectamine 3000 (ThermoFisher Scientific) following the manufacturer’s instruction. For a well in a 6-well plate, typically ∼1 × 10^6^ cells were seeded and transfected with 2.5 μg of DNA.

### rDNA barcoding

The K562 PEmax(+) cells were used in the barcoding experiment. For each round of barcoding, 2 × 10^5^ cells were nucleofected with 500 ng the pegRNA library and 500 ng pmax-GFP (Lonza). Cells were passaged 2 days after nucleofection. On day 4, a portion of cells were harvested for gDNA and amplicon sequencing, while the rest were used for the next round of nucleofection. A total of 10 rounds of barcoding were performed on the cells. Single clone derivation happened 10 days after the last nucleofection using a limited dilution method.

### Polysome fractionation

Polysome profiling of K562 cells was performed as described in Yale Cooperative Center of Excellence in Hematology (YCCEH) protocol *Polysome Preparation from Eukaryotic Cells*, contributed by Manoj Pillai Lab. In brief, the TMK-100 lysis buffer was prepared in nuclease-free water and contained 10 mM Tris-HCl (pH 7.4), 5 mM MgCl₂, 100 mM KCl, 1% (v/v) Triton X-100, and 0.5% (w/v) sodium deoxycholate. Immediately before use, the buffer was supplemented with 1 U/mL RNase inhibitor and 2 mM dithiothreitol (DTT). Sucrose gradient solutions containing 10%, 20%, 30%, 40%, 50%, 60% (w/v) sucrose were prepared in nuclease-free water with 100 mM KCl, 5 mM MgCl₂, 2 mM DTT, and 20 mM HEPES-KOH (pH 7.4). Linear sucrose gradients were prepared manually by sequentially layering sucrose solutions (10%, 20%, 30%, 40%, 50%, 60%) into a tube for the SW41Ti rotor (Beckman) using a syringe fitted with a long blunt-ended needle, as described by Sobhany et al. (*62*). The layered gradients were flash frozen at -80°C until further usage. The day before harvesting the cells, gradients were kept at 4°C overnight to let them settle. 4 × 10^7^ K562 cells were harvested and lysed on ice for 10 min. The lysate was centrifuged at 21,000 ×g for 10 min at 4 °C and the supernatant was loaded onto the sucrose gradient. The tube was centrifuged at 222,228 × g in a Beckman Coulter ultracentrifuge for 2 hours at 4 °C. After that, the gradient was manually fractionated from top to bottom. Absorbance at 260 nm was obtained. Fractions belonging to free RNA, monosome, and polysome were pooled and subjected to RNA isolation with TRIZol LS (Thermofisher Scientific).

### Amplicon sequencing

Cells were lysed in a lysis buffer (100 mM Tris-HCl pH8.0, 0.05% SDS and 0.04 mg/mL proteinase K), and incubated at 50 °C 60 min and 80 °C 30 min. Total RNA was extracted with TRIZol LS and reverse transcribed using SuperScript IV (Thermofisher Scientific) following manufacturers’ instructions. gDNA lysates, cDNA, and plasmid samples were directly used for PCR with the KAPA2G Robust HotStart ReadyMix with primers (0.5 μM each) designed for rDNA or the pegRNA construct. A touchdown PCR reaction was performed as follows: 95 °C 3 min; 95 °C 15 s, 66-N °C 15 s, 72 °C 40 s, for 8 cycles (N was the real-time cycle number); 95 °C 15 s, 60 °C 15 s, 72 °C 40 s, for M cycles (decided by qPCR); and 72 °C 1 min. The PCR product was purified with AMPure XP beads. 5 ng PCR product was re-amplified with the Nextera P5 (p54) and TruSeq P7 (p55) index PCR primers in **table S6** for 5 cycles. The final libraries were cleaned up with AMPure XP beads and sequenced on an Illumina NextSeq 2000 sequencer.

Sequencing reads were demultiplexed using the bcl2fastq software (Illumina). A custom script was used to determine editing rates and barcode abundance.

### Chromosome spreads, Fluorescent In-Situ Hybridization (FISH)

The chromosome spreads and FISH experiment were performed following instructions in Potapova et al., 2025 (*18*). For the preparation of chromosome spreads, cells were blocked in mitosis by the addition of Karyomax colcemid solution (0.1 µg/ml, Thermofisher Scientific) for 6-7 h and collected by spinning down at 250 ×g for 5 min. Collected cells were resuspended in a hypotonic 0.4% KCl solution, incubated for 10 min and pre-fixed by addition of methanol:acetic acid (3:1) fixative solution (1% total volume). Pre-fixed cells were spun down and resuspended in methanol:acetic acid fixative. Spreads were dropped on a glass slide and incubated at 65 °C overnight. Before hybridization, slides were treated with 0.1 mg/mL RNAse A (Qiagen) in 2× SSC for 45 min at 37 °C and dehydrated in a 70%, 80%, and 100% ethanol series for 2 min each. Slides were denatured in 70% deionized formamide/2× SSC solution pre-heated to 72 °C for 1-2 min. Denaturation was stopped by immersing slides in 70%, 80%, and 100% ethanol series chilled to -20 °C. Labeled DNA probes were denatured separately in a hybridization buffer by heating to 80 °C for 7 min before applying to denatured slides. Fluorescently labeled rDNA BAC probe RP11-450E20 was from Empire Genomics. CenSat D14Z1/D22Z1 labeled probe was from Cytocell (LPE 014). Whole chromosome paints for acrocentric chromosomes were from Applied Spectral Imaging (FPRPR0167).

Specimens were hybridized to the probes under a HybriSlip hybridization cover (GRACE Biolabs), sealed with Cytobond (SciGene) and incubated in a humidified chamber at 37 °C for 48-72 hrs. After hybridization, slides were washed in 50% formamide/2× SSC 3 times for 5 min per wash at 45 °C, then in 1× SSC solution at 45 °C for 5 min twice, and at room temperature once. For biotin detection, slides were incubated with streptavidin conjugated to Cy5 (Thermo Fisher) for 2-3 hrs in PBS containing 0.1% Triton X-100 and 5% bovine serum albumin (BSA), and then washed 3 times for 5 min with PBS/0.1% Triton X-100. 1 µg/mL DAPI was added to final washes to label chromosomes. Slides were mounted in ProLong Gold or ProLong Glass (Thermo Fisher). Confocal Z-stack images were acquired on the Nikon TiE microscope equipped with PlanApo 100x oil immersion objective NA 1.45, Yokogawa CSU-W1 spinning disk, ORCA-Fusion BT sCMOS camera (Hamamatsu), and NIS Elements software. All image processing was performed using ImageJ/FIJI. Maximum intensity projections of confocal Z-planes were generated, and rDNA-containing chromosomes segmented, identified and arranged into karyograms.

### ONT ultra-long sequencing library preparation

For each library, 6 × 10^6^ K562 cells were harvested and high-quality gDNA was prepared using the Monarch HMW DNA Extraction Kit (NEB, T3060). The sequencing library was prepared using the Ultra-Long DNA Sequencing Kit V14 (ONT, SQK-ULK114) and loaded onto a PromethION flowcell. Basecall was performed using Dorado.

### Barcode-guided NOR assembly

17 ONT ultra-long-read sequencing runs were performed on the 28S rDNA barcoded clone 6. A total of 900 Gb of sequencing data were generated. To assemble the NORs, a barcode-guided approach was taken (**fig. S2A**).

#### 1. ONT reads preparation

As the barcodes were inserted at the Cas9 cut site by design, ONT reads containing the first 17 bases of the spacer sequence (TGGCGGAGATGGGCGCC) or its reverse complement form (GGCGCCCATCTCCGCCA) were extracted from the BAM files (n = 15,273). Sequencing data from 14 out of 17 runs were base-called with alignment, resulting in a subset of reads being reported multiple times as supplementary alignments (1,562 supplementary alignments contained within 1,149 unique reads). Therefore, 13,711 unique reads were obtained after deduplication. Reads containing only the reverse sequence were reverse-complemented. Reads containing both forward and reverse spacer sequences were manually inspected (n = 19), among which 7 reads were confirmed to be palindromic, with 6 containing 2 barcodes, and 1 containing 4 barcodes. Although these reads were not used for assembly, their barcode identity and order were later confirmed to support our assembly results. The remaining 12 were found to contain local inversions, and were retained for subsequent analyses, later summarized in Type 5-3 (**fig. S4E**).

After removing the palindromic reads (n = 7), barcodes were extracted by ambiguous matches with a left anchor sequence (TGGCGGAGATGGGCGCC, allowing up to two mismatches) and a right anchor sequence (GCGAGG, allowing one mismatch). Barcode sequences ranging from 6 to 30 bp were retained (n = 16,317). Low-abundance barcodes in reads containing at least two barcodes (count ≤ 25, n = 213) were subjected to contextual correction. Correct barcode sequences were inferred from neighboring barcodes in all other reads (n = 185). In addition, barcode positions and inter-barcode distances in reads containing at least two barcodes were carefully examined to recover missing barcodes caused by sequencing errors in the anchor sequences or unedited barcoding sites (represented as “WT” later in the assembly). Based on the corrected barcode assignments, these remaining reads were divided into three groups: reads containing a single barcode (n = 11,571), reads containing two barcodes including “WT” (n = 1,425), and reads containing at least three barcodes including “WT” (n = 708).

#### 2. Flank sequence identification

To identify reads spanning NOR boundaries, we searched each read for regions matching the rDNA unit reference (KY962518.1) (*4*) using exact 20-mer matches. Regions containing fewer than 20 consecutive matching 20-mers were treated as mismatches. A region was considered a true rDNA-matching region when at least 300 matching 20-mers were detected within a 1-kb window. The unmatched sequences extending from the read ends to the boundaries of the rDNA-matching region were designated as left or right flanks accordingly. Flanks of at least 25 kb were retained for further analysis.

Among the single-barcode reads, 418 flank-containing reads were identified (left flanks, n = 291; right flanks, n = 127). Among reads containing at least two barcodes, 189 flank-containing reads were identified (left flanks, n = 106; right flanks, n = 80; both left and right flanks, n = 3). The flank sequences identified from reads containing at least two barcodes were used to collapse redundant information and define unique flank sequences. Single-barcode reads were then used to validate the identified flanks. After inspecting the read distribution of the unique flank information (**fig. S2C**), we excluded flanks supported by only one read. This yielded 18 left flanks and 18 right flanks with minimal 4 supporting reads (**table. S2**), which were used for barcode scaffold assembly.

#### 3. Barcode-only scaffolds construction

Reads containing at least 3 barcodes were used for scaffold construction (n = 708). Barcode order (paths) within individual reads were extracted and merged using an overlap-based assembly strategy. Barcode paths were ranked by length, and pairs of paths sharing at least two consecutive barcodes were merged according to their maximal suffix–prefix overlap. When multiple merging configurations occur, the junction was resolved by inspecting inter-barcode distances across reads. For each shared barcode pair, observed inter-barcode distances were grouped into clusters using one-dimensional single-linkage clustering. A candidate join was accepted only when the shared barcode pairs belonged to the same distance cluster. After each new merge, any barcode path that was completely contained within the newly generated longer path was removed to eliminate redundant assemblies. The merging process was performed iteratively until no additional overlaps could be identified, resulting in a set of maximally extended barcode scaffolds. During this process, 11 reads were excluded, among which 3 reads contained multiple low-occurrence barcodes that could not be reliably corrected, whereas the remaining 8 reads contained alternative path junctions supported by only a single read, in contrast to 5-10 supporting reads for the corresponding major paths. This procedure yielded 16 barcode paths spanning from the left flank to the right flank of the actual NORs. Four barcode paths (two left-flank-containing paths and two right-flank-containing paths) could not be joined based on two-barcode overlaps; however, fortunately, the two pairs could be confidently linked using unique barcodes spanning their respective junctions. Together, these steps resulted in 18 high-confidence scaffolds, each being anchored by a left and a right NOR flanking sequence, and being supported by a median of 12-24 independent reads (**Fig. 2B**). These scaffolds were therefore retained as the framework for NOR assembly.

#### 4. Alignment and consensus sequence construction

All remaining reads containing at least 2 barcodes were subsequently aligned to the barcode-only scaffold according to the barcode order (n = 2,122). If a read could be aligned to multiple positions based on barcode match, inter-barcode distances were calculated in both the read and the existing barcode paths, and clustered. The read was assigned to the path with the best-matching inter-barcode distances. Remaining ambiguities were resolved using BLASTN from the BLAST suite (v2.17.0) (*63*), based on sequence similarity to reads assigned to the candidate positions. After alignment, 29 reads that could not be unambiguously assigned to the scaffolds were discarded, including 17 reads containing low-occurrence barcodes that could not be reliably corrected. Each aligned read was split into barcode-defined fragments, including inter-barcode fragments located between consecutive barcodes, and terminal fragments located before the first barcode or after the last barcode. Each fragment was assigned to a group corresponding to its position on the barcode-only scaffold. For each inter-barcode position on the scaffold, the mean length of the corresponding intact inter-barcode fragments was calculated across all supporting reads in the group. A read was removed if any of its inter-barcode fragments deviated by more than 2.5% from the mean length of the corresponding inter-barcode group, or if either terminal fragment was 2.5% longer than the mean length. This filtering step removed 32 reads. Fragments from each group, derived from the remaining 2,061 reads, were then aligned separately using MAFFT (v.7.526) with the “--nofft” option to perform alignment without the FFT-based initial alignment step (*64*), and consensus sequences were generated based on the most frequent nucleotide (or gap) at each aligned position. For each position, the effective coverage (number of valid sequences spanning the position) and the number of sequences supporting the consensus state were calculated. A position-specific consensus score was then defined as the ratio between supporting reads and all reads (**fig. S2D**). The interval consensus sequences were then concatenated according to barcode order to generate the NOR assemblies **(data S1**).

#### 5. Manual curation of the assemblies

The consensus sequences of the rDNA units were standardized by defining the unit start as sequence “AAAATGAAAATGAAAATGAAACGCAACAAAATAATTAAAAAGTGAGTTTCTGGGGAAAAA”. This boundary was selected as it marks the left most end of the repetitive sequences in 15/18 NORs. 20-mers immediately downstream of this sequence were repetitive while 20-mers upstream were unique, indicating a transition into rDNA units. Consequently, the unit end was defined using the sequence “CCAGCCTGGGCGACAGAGCGAGACTCCGTCTCCAA”, which is upstream of the start sequence in the NORs. 3 NORs (NORs 1, 11, 12) had truncations in their first units and therefore lacked the defined unit start sequence. Terminal units in 15/18 NORs were truncated and extended only to either 18S or ITS1. These truncation events have been observed in the CHM13 and HG002 genomes (*10*, *11*). In addition, the remaining 3 NORs (NORs 2, 7, 18) exhibited 2 distinct truncation breakpoints (**table S3**).

After this curation, a total of 290 rDNA units were identified across all assembled NORs. The first truncated units of NORs 11 and 12 were retained as the missing portions only accounted for less than 10% of the units. In contrast, the first unit of NOR1 and all the 18 trailing truncated units were excluded from downstream analysis (such as sequence variation analysis). In summary, 271 rDNA units were carried forward, among which 265 contained identifiable barcode sequences, whereas 6 units did not contain a barcode or lacked an intact barcoding site.

### Pairwise similarity analysis of rDNA units

Pairwise sequence similarity between rDNA units was calculated using an all-versus-all alignment approach with Minimap2 (V2.30, Oxford Nanopore overlap mode, -x ava-ont) (*65*, *66*). For each comparison, the highest-scoring alignment was retained to avoid redundancy from multiple local alignments. Sequence similarity was calculated as the ratio between the number of matched bases and the total alignment length. To compare the differences between NORs, unit pairs were grouped as intra-NOR and inter-NOR and visualized as a cumulative distribution function (CDF) plot (**Fig. 2E**). *P*-value was calculated using the Wilcoxon rank-sum test.

### Annotation and structural visualization of rDNA units

To characterize the structural variations among individual rDNA units, each unit was aligned against the reference sequence (KY962518.1) (*4*) using a hash table–based exact k-mer mapping approach (*67*). The k-mer size was selected as the smallest value for which more than 99% k-mers in the reference sequence were unique. This resulted in a k-mer size of 150 bp. A reference 150-mer index was constructed for the reference sequence, and query sequences were scanned for exact matches to these 150-mers. Both forward and reverse-complement matches were considered. Matched 150-mers were used to establish a positional correspondence between each query unit and the reference sequence.

For annotation, the main features found in the reference sequence were lifted onto the consensus sequences based on 20-mer mapping status. Features shorter than 80 bp were excluded to reduce spurious assignments. Adjacent fragments annotated with the same feature type were merged. The final annotations were exported into GFF3 format for downstream analyses.

Structural variations were visualized by comparing the positional correspondence between query and reference sequences. Regions showing consistent coordinate offsets were merged into continuous structural blocks, whereas abrupt changes in offset values were interpreted as candidate structural rearrangements. For each unit, a dot-plot–style visualization was generated by plotting query coordinates against the reference coordinates. Discontinuous mapping segments were highlighted to facilitate the identification of insertions, deletions, and inversions (**data S2**).

### Principal component analysis (PCA)

To investigate sequence variations among rDNA units, each of the 271 units was represented by its k-mer composition. To determine the optimal k-mer length, k values ranging from 2 to 10 were evaluated based on PCA performance (variance captured) and clustering quality (silhouette score). For each tested k value, k-mer frequencies were calculated across the entire unit sequence and normalized by the total k-mer count. The resulting k-mer frequency matrices were standardized using z-score normalization and subjected to PCA. A k-mer size of 6 was selected because it provided a clean separation of unit groups by *k*-means clustering (*k* = 6), while avoiding the low sequence specificity associated with smaller k values and the increased feature sparsity observed for larger k values (**fig. S4A**). To group similar units, *k*-means clustering (*k* = 6) was performed on the PCA-transformed data. The resulting cluster distributions were visualized in the PC1-PC2 space. As the largest cluster (Cluster 1) contained 201 of the 271 units, a second PCA was performed to resolve finer differences within the group. The PCA was performed based on 3-mer frequencies. *k*-means clustering (*k* = 9) was applied to group the units. Consensus sequences of clusters or subclusters were generated by aligning the consensus sequences of units within that cluster or subcluster using MAFFT with the “--nofft” option, based on the most frequent nucleotide (or gap) at each aligned position.

To enable comparisons with other human reference genomes, we examined the rDNA sequences of the CHM13 (*10*) and HG002 (v1.1) (*11*). In HG002, only the terminal portions of the rDNA arrays are assembled, with the intervening regions represented by “N”s in the reference genome. In CHM13, the intervening rDNA units are represented by exact duplications of the assembled terminal unit sequences. We therefore extracted the assembled terminal rDNA units from HG002, while for CHM13, we collapsed the exact-copy units into unique representative sequences. The resulting sequences were encoded using the same k-mer frequency scheme and projected onto the previous PCA plots.

### Classification of rDNA units

2 rounds of PCA and clustering classified the 271 rDNA units into 14 groups. To ensure accuracy of the grouping, all units were aligned to the reference rDNA sequence (KY962518.1) using 150-mer matches and visualized (**data S2**). Units that displayed inconsistent structural patterns with the majority of the group members were examined and re-grouped. This manual curation resulted in 25 groups (**Fig. 3C**; **fig. S4E**; and **table S4**).

Based on features and regions involved in SVs, the 25 groups were organized into 5 primary types: Type 1 represents units with structures most similar to the reference sequence (KY962518.1); Type 2 includes units with SVs affecting the R repeats in the IGS; Type 3 comprises units with SVs in the Butterfly/Long repeats in the IGS; Type 4 contains units harboring SVs in both R repeats and Butterfly/Long repeats; Type 5 consists of units with complex rearrangements affecting the 47S pre-rRNA region. Within each primary type, distinct groups were designated as “subtypes”. The subtypes were ordered in descending order based on the number of units they contained.

Consensus sequences of subtypes were generated by aligning the consensus sequences of units within that subtype using MAFFT with the “--nofft” option, based on the most frequent nucleotide (or gap) at each aligned position.

### *De novo* genome assembly and chromosomal assignment of NOR-containing scaffolds

*De novo* assembly of the K562 genome was performed using hifiasm (v0.25.0-r726) in ONT mode (*56*, *57*, *68*). Prior to assembly, reads shorter than 100 kb were removed using SeqKit (v2.13.0) (*69*). The previously assembled NOR sequences were then converted to FASTQ format with uniformly high-quality scores and appended to the filtered ONT read data. This yielded 119 contigs (N50 = 99,789,102; **table S1**). Primary contigs were extracted from the resulting assembly graph using gfatools (v0.4-r214; https://github.com/lh3/gfatools) and aligned to the CHM13 reference genome using minimap2. The alignment results were used both to annotate the assembled contigs and to connect primary contigs into chromosome-scale scaffolds, with scaffold organization guided by the corresponding FISH results. Because the short arms of the acrocentric chromosomes exhibit high sequence similarity, alignments within these regions were often fragmented and ambiguously assigned. Therefore, alignment fragments shorter than 1 Mb that mapped to different acrocentric chromosomes were considered ambiguous and were not used for chromosome assignment. These ambiguous fragments were displayed in grey in the scaffold annotation (**Fig. 5**).

To determine the chromosomal locations of the NORs, NOR flanking sequences were aligned to the assembled scaffolds using minimap2. Because alignments to the short arms of acrocentric chromosomes were often difficult to differentiate, we first used unambiguous flank alignments from translocated NORs to establish their locations. This assigned NOR1 to der(21)t(1;21), NOR2 to chr9, and NORs 11 and 12 to der(17)t(9;17). We then considered high-confidence alignments (MAPQ > 10), assigning NORs 5 and 6 to chr14, NORs 8-10 to chr15, and NORs 13 and 14 to chr21. After relaxing the MAPQ threshold, among the remaining NORs, only NOR4 was aligned to chr13; NORs 17 and 18 were aligned to the short and long arm of the Philadelphia chromosome, respectively; while NORs 15 and 16 were aligned to the short arms of the two normal chr22, thereby establishing the locations of these NORs. The remaining two NORs (NORs 3 and 7) showed substantial length differences between each other, and were therefore assigned to the remaining NOR-bearing chromosomes (der(13)t(9;13) and chr14) based on their relative FISH signal intensities. For NORs mapped to duplicated chromosomes, the final assignment was guided by rDNA signal intensity in FISH (**fig. S3**). As a sanity check, we compared the sequence similarity between NOR flanks using BLASTN in the BLAST suite. The sequence similarity between flanks assigned to duplicated chromosomes was higher than that between flanks assigned to different chromosomes.

Assembly completeness of the original output of hifiasm was evaluated using BUSCO (v6.0.0) (*70*) with the primates_odb12 lineage dataset. Accuracy of the final assembly of the NOR-containing scaffolds was evaluated using Merqury (v1.3) (*71*) based on k-mer spectra derived from the original ONT sequencing reads.

### Loss of heterozygosity (LOH) analysis

LOH analysis was conducted based on the general framework described previously (*72*), with modifications appropriate for this study. Single-nucleotide polymorphisms (SNPs) were identified using Clair3 (v1.0.10) (*73*). Variants with sequencing depth ≥ 30× and quality score ≥ 20 were retained for downstream analyses. For each SNP, the variant allele frequency (VAF) was calculated as the proportion of reads supporting the alternative allele relative to the total read depth (**fig. S6B**). Genome-wide LOH assessment was performed in non-overlapping 500-kb windows. Within each window, SNPs were classified according to their VAF values as AB-like (0.4-0.6) or AAB-like (0.25-0.4 or 0.6-0.75), representing diploid-like or trisomy-like allele configurations, respectively. For each window, the proportions of AB-like and AAB-like SNPs were calculated. Windows containing >30% AB-like SNPs were classified as AB, whereas those containing >30% AAB-like SNPs were classified as AAB. Windows not meeting either criterion were classified as LOH.

### Pairwise similarity analysis of NORs

The similarity between the NORs was evaluated using two complementary approaches.

The first approach was based on unit level sequence similarity, in which pairwise sequence comparison among individual rDNA units were first calculated. To obtain NOR-level similarity estimates, a reciprocal best-hit (RBH) strategy was employed. For each pair of NORs, the highest similarity scores between every unit in one NOR and all units in the other NORs were determined, and the mean of these best-hit similarities was calculated. The procedure was performed in a reciprocal way, and the average scores were designated as the final similarity scores for all the NOR pairs. Self-similarity scores were set to 1.0. For plotting, NORs were clustered on the similarity matrix by agglomerative hierarchical clustering (Ward’s minimum-variance method, Euclidean distance) and displayed as a heatmap (**Fig. 4B**).

The second approach assessed the similarity directly at the NOR level. A k-mer-based approach was implemented using Mash (Version 2.3) (*74*). MinHash sketches were generated for each NOR using a k-mer size of 31 and a sketch size of 100,000. Pairwise Mash distances were calculated in an all-versus-all manner and converted to Jaccard similarity coefficients using the inverse Mash distance transformation. The resulting similarity matrix was subsequently normalized using the Sinkhorn-Knopp algorithm to obtain a doubly stochastic matrix (*75*), which was used for downstream clustering and visualization (**fig. S6A**).

### Methylation analysis

To characterize the DNA methylation patterns across NORs, reads used for consensus sequence construction (n = 2,061) were split into fragments according to the barcode-defined intervals, and each fragment was aligned to the previously defined consensus sequence of the interval using minimap2 (-ax map-pb). This interval-based alignment strategy minimized potential mis-alignment caused by reads spanning multiple rDNA units. Alignment coordinates, CIGAR strings, and read orientation information were extracted from the resulting SAM files. For each ONT read, the CIGAR strings obtained from individual fragment alignments were concatenated according to the original fragment order within the read, generating a single alignment representation spanning the assembled NOR sequence. The reconstructed CIGAR strings were then substituted into the SAM files derived from the Dorado output, thereby adding the alignment information while preserving the original methylation annotations. Reads aligned in the reverse-complement orientation were assigned a SAM flag of 16, whereas forward-aligning reads were assigned a flag of 0.

DNA methylation signals were subsequently extracted from the processed SAM file using the Modkit in pileup mode (v0.6.1; https://github.com/nanoporetech/modkit). The 5-methylcytosine (5mC) calls were filtered using the estimated pass threshold automatically determined by Modkit (0.53125). Regional methylation levels were calculated in 50-bp windows across the assembled NOR sequences using the “stat” module of Modkit as the proportion of modified bases in all modification-callable (CpG) sites within each window. For each annotated rDNA feature, the feature-level methylation was calculated as the total number of modified bases divided by the total number of modification-callable bases.

To determine which rDNA genomic feature best distinguished the high-methylated and low-methylated units, DNA methylation levels were calculated for each annotated feature or combinations of the features across all complete rDNA units. For each feature independently, *k*-means clustering (*k* = 2) was performed using the per-unit methylation levels, and clustering performance was evaluated using the silhouette score, with higher scores indicating stronger separation between the two methylation states. As the 47S pre-rRNA region, along with the upstream Pol I enhancer and promoter, constitutes the transcriptional cassette of the rDNA unit and exhibited the highest silhouette score among all annotated features, unit methylation status was defined based on methylation levels calculated within this region. Therefore, units that were lowly methylated in the 47S region were considered “low methylation”, while units with heavy methylation in this region were classified as “high methylation” (**Fig. 2A**; **Fig. 6A**). The 5mC levels were summarized as genomic tracks for visualization (**data S3**).

### Barcode abundance estimate in short-read sequencing

The sequence surrounding the rDNA barcodes were amplified (**table S6**) and subjected to amplicon sequencing (Illumina). For the 28S barcoded clone, barcode abundance was measured in both gDNA and total RNA samples (3 replicates each). Barcode sequences were extracted from Read 1 (antisense) by searching for pattern CCNNNN[Barcode]GGCGCCCATCTCCGCCA, where the barcode length could range from 4 to 8 bp. Extracted sequences were reverse-complemented to match the barcode orientation in the NOR assemblies. Abundance of barcodes present in the NOR assemblies was calculated separately for rDNA and rRNA libraries, averaged across three technical replicates.

### Integrated visualization of NORs

To summarize the genetic and epigenetic landscape of the assembled NORs, multiple layers of information, including the general NOR architecture, structural variations, long-read alignment status, DNA methylation, transcriptional activity, and rDNA unit classifications, were integrated into a ring plot for easy visualization, using the circlize package (v.0.4.17) in R (v.4.2.1) (**Fig. 2A**).

At a high level, the NORs were ordered based on their putative chromosomal origins and sequence similarity. For structural variation, insertions, deletions, and inversions larger than 500 bp were highlighted. ONT reads > 150 kb were selected and aligned to their corresponding supporting NORs. DNA methylation levels were calculated by intersecting modkit-derived methylation profiles with the 47S pre-rRNA, and the upstream Pol I enhancer and promoter region. Two units in which more than 60% of the 47S pre-rRNA region was missing (Type 5-2 and 5-3) are marked grey at corresponding positions (“not available”). The transcriptional activity measured by short-read amplicon sequencing was plotted for units with unique barcodes. All other units are marked in grey (“not available”).

## SUPPLEMENTARY FIGURES

**fig. S1.**
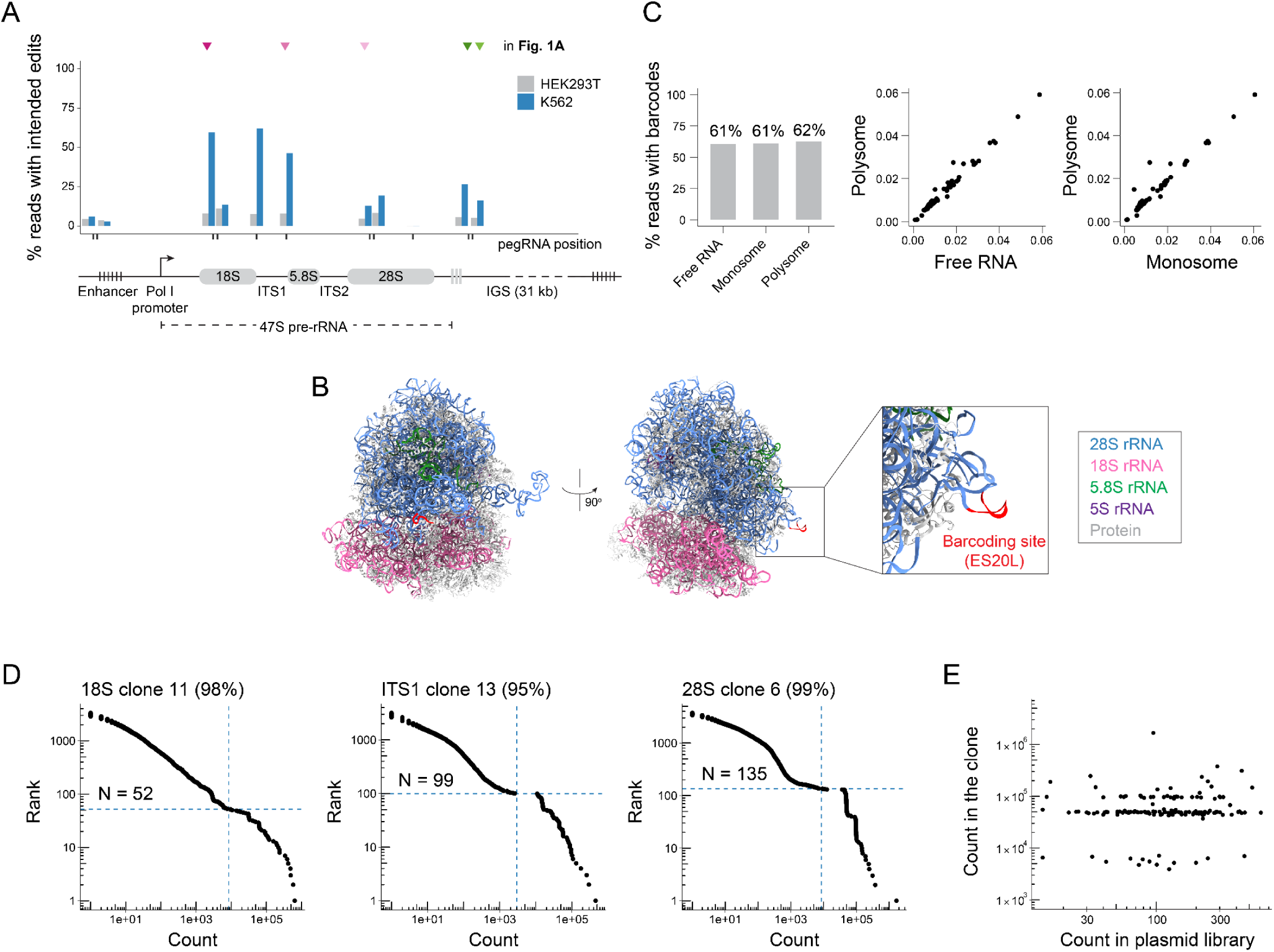
Prime editing-mediated rDNA barcoding. **(A)** Editing efficiencies of 11 pegRNAs targeting rDNA in K562 and HEK293T cells. Five pegRNAs (marked by triangles) were selected for comparison with the HEK3.insCTT pegRNA. **(B)** Cryo-EM structure of a human ribosome (PDB:4UG0). The barcoding site on ES20L is highlighted in red. **(C)** Polysome profiling of a partially barcoded monoclonal line (62% barcoding rate in gDNA). Percent reads with barcodes in free RNA, monosome, and polysome fractions are compared. **(D)** Barcode read count and rank in the monoclonal lines with the highest barcoding rates in each barcoding experiment. Estimated barcode numbers are annotated. x- and y-axes in log_10_ scale. **(E)** Scatter plot for barcode read count in the 28S clone 6 and in the original plasmid library. The lack of correlation between the two indicates the occurrence of duplicated barcodes can not be attributed to biased barcode representation in the plasmid library. x- and y-axes in log_10_ scale.

**fig. S2.**
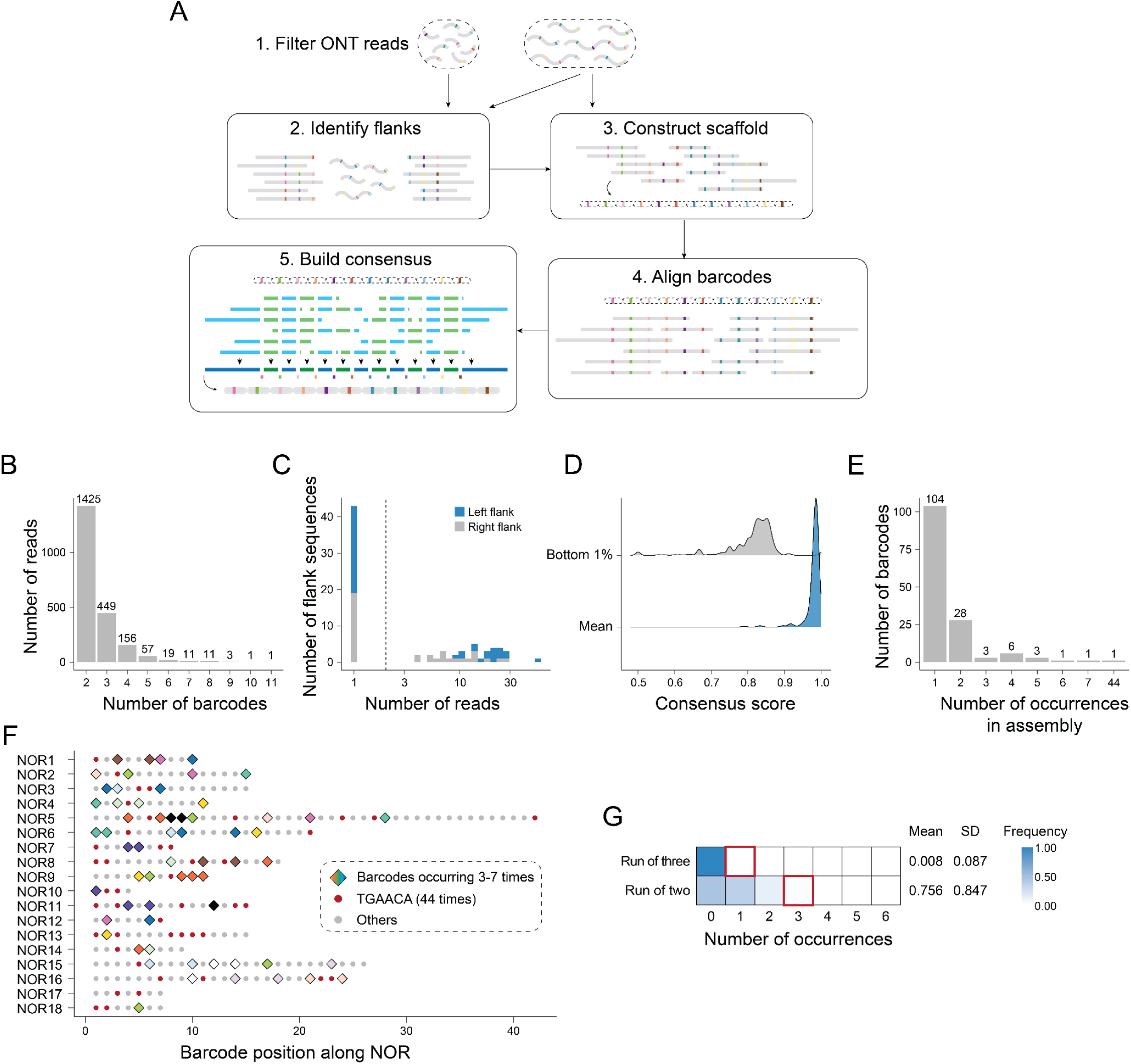
Barcode-guided NOR assembly. **(A)** Schematic of the barcode-guided NOR assembly approach (see **Methods**). **(B)** Number of rDNA barcodes in the ONT reads containing at least two barcodes (n = 2,133). **(C)** Number of reads supporting each flank sequence. Flanks supported by at least 4 reads (on the right of the dashed line) were used for assembly. X-axis in log_10_ scale. **(D)** Positional consensus scores were calculated for all barcode intervals (see **Methods**). Distributions of mean scores and scores at the 1st percentile positions are plotted separately. **(E)** Number of occurrences of each barcode in the assembly. **(F)** Structure of the 18 barcode-only scaffolds. 265 barcode positions are shown. Barcodes occurring 3-7 times are shown as filled diamonds and colored. The 44 “TGAACA” are red circles. All other barcodes are grey. **(G)** Frequencies of “runs of two” and “runs of three” consecutive occurrences of the same barcode observed across 10,000 permutation experiments. The 14 barcodes occurring 3-7 times were used for this simulation. The mean number of occurrences and the standard deviation are annotated. Red squares indicate the observed number of occurrences under each condition.

**fig. S3.**
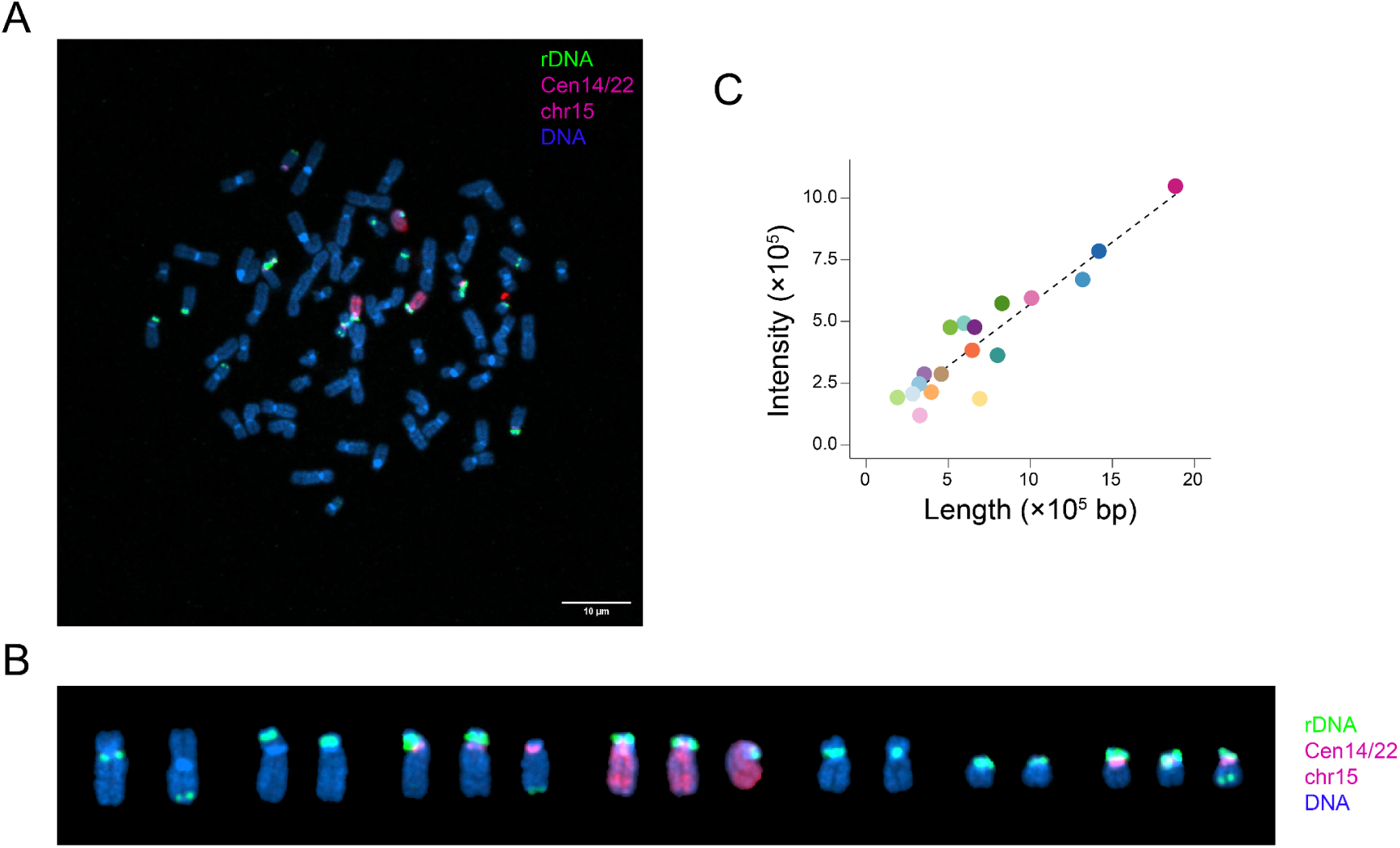
FISH analysis of the K562 rDNA-barcoded clone. **(A)** A representative image of the chromosome spreads and FISH result of the K562 barcoded clone. rDNA (green), the centromeres of chrs. 14 and 22 (red), and chr 15 (red) are stained with probes. DNA is stained with DAPI. **(B)** NOR-containing chromosomes in **A** are isolated and arranged. **(C)** Scatterplot showing correlation between the FISH rDNA signal and the lengths of corresponding NORs. The 18 NORs are colored as in **Fig. 2**.

**fig. S4.**
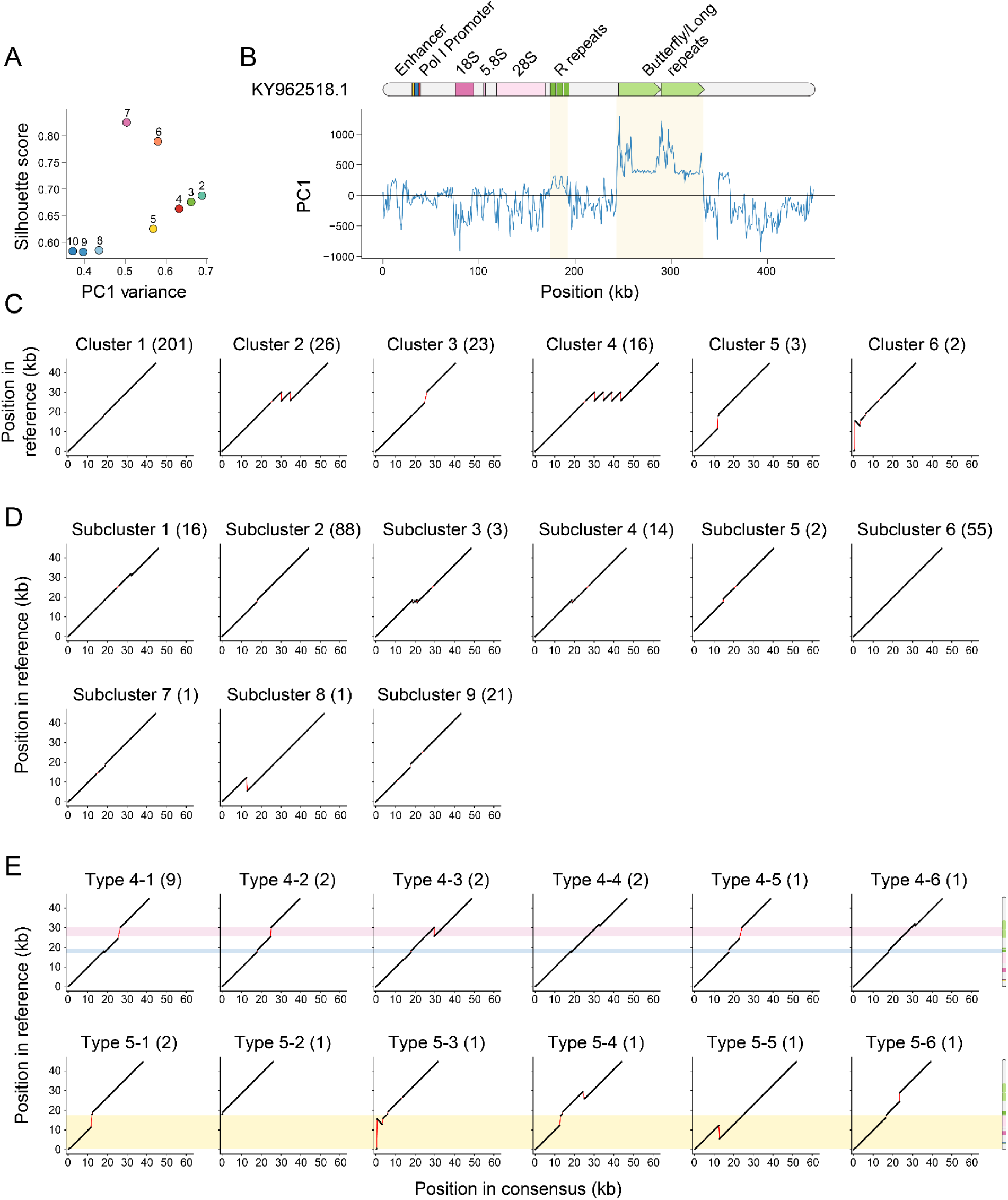
K-mer-based classification of rDNA units. **(A)** Scatter plot showing the relationship between the proportion of variance explained by the first principal component (PC1; x-axis) and the clustering quality of a *k*-means clustering (*k* = 6; silhouette score; y-axis) across different k-mer sizes. **(B)** PC1 values of consecutive 100-bp bins along the rDNA reference sequence (KY962518.1), obtained by projecting the 6-mer composition of each bin onto the PCA model shown in **Fig. 3A**. **(C)** Consensus sequences of units belonging to the 6 main clusters are aligned to the reference allele KY962518.1. **(D)** Consensus sequences of units belonging to the 9 subclusters are aligned to the reference allele. **(E)** Consensus sequences of unit groups in Type 4 and 5 are aligned to the reference allele. Subtypes are numbered in descending order of unit count. Altered regions are highlighted and project onto a schematic of the reference allele at right.

**fig. S5.**
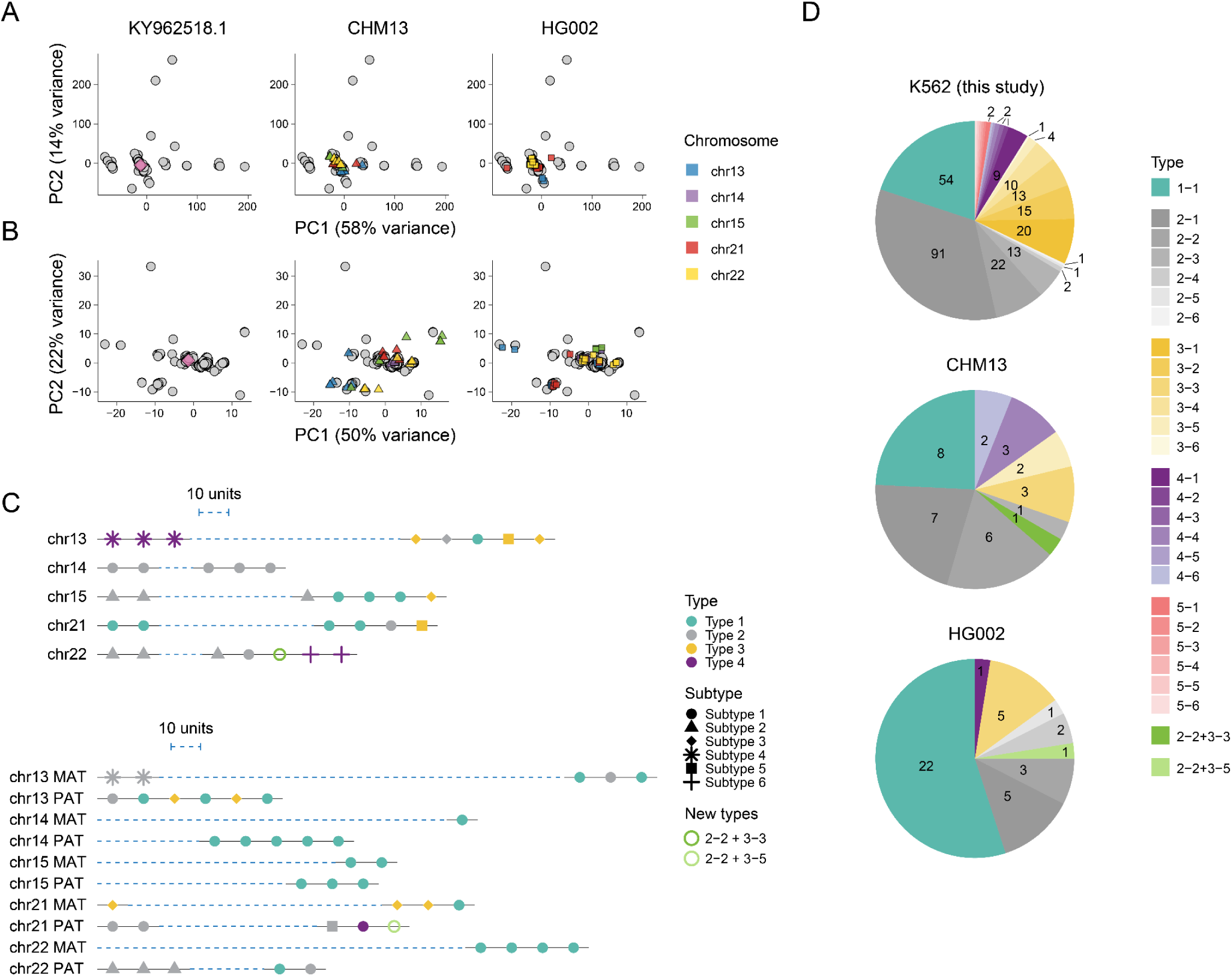
Comparison of rDNA units in K562 cells with those from standard genomes. **(A)** PCA of the 271 rDNA units based on 6-mer content, showing PC1 and PC2 (**Fig. 3A**). The reference rDNA allele KY962518.1 (left), and rDNA units from CHM13 (middle) and HG002 (right) are projected onto the PCA space. rDNA units are colored by chromosomal origin. (**B**) PCA of the 201 cluster 1 units based on 3-mer content, showing PC1 and PC2 (**Fig. 3B**). The reference rDNA allele KY962518.1 (left), and rDNA units from CHM13 (middle) and HG002 (right) are projected and colored by chromosomal origin. **(C)** Schematic architecture of the NORs obtained from the standard genomes: CHM13 (top) and HG002 (bottom). Only rDNA units close to NOR ends were used for comparison (same units as in **A-B**). The units were classified according to **Fig. 3C** and **fig. S4E**. Most rDNA units fell into known groups, except for two units found to comprise SV features of Type 2-2 and 3-3 (CHM13), and Type 2-2 and 3-5 (HG002), respectively. Dashed blue lines represent compressed sequences between extracted rDNA units. The lengths of these lines correspond to estimated NOR sizes in their respective data sources. The scale bar represents the length of 10 rDNA units. **(D)** Pie charts showing rDNA type composition across genomes. The new types in CHM13 (Type 2-2 + 3-3) and HG002 (Type 2-2 + 3-5) are green.

**fig. S6.**
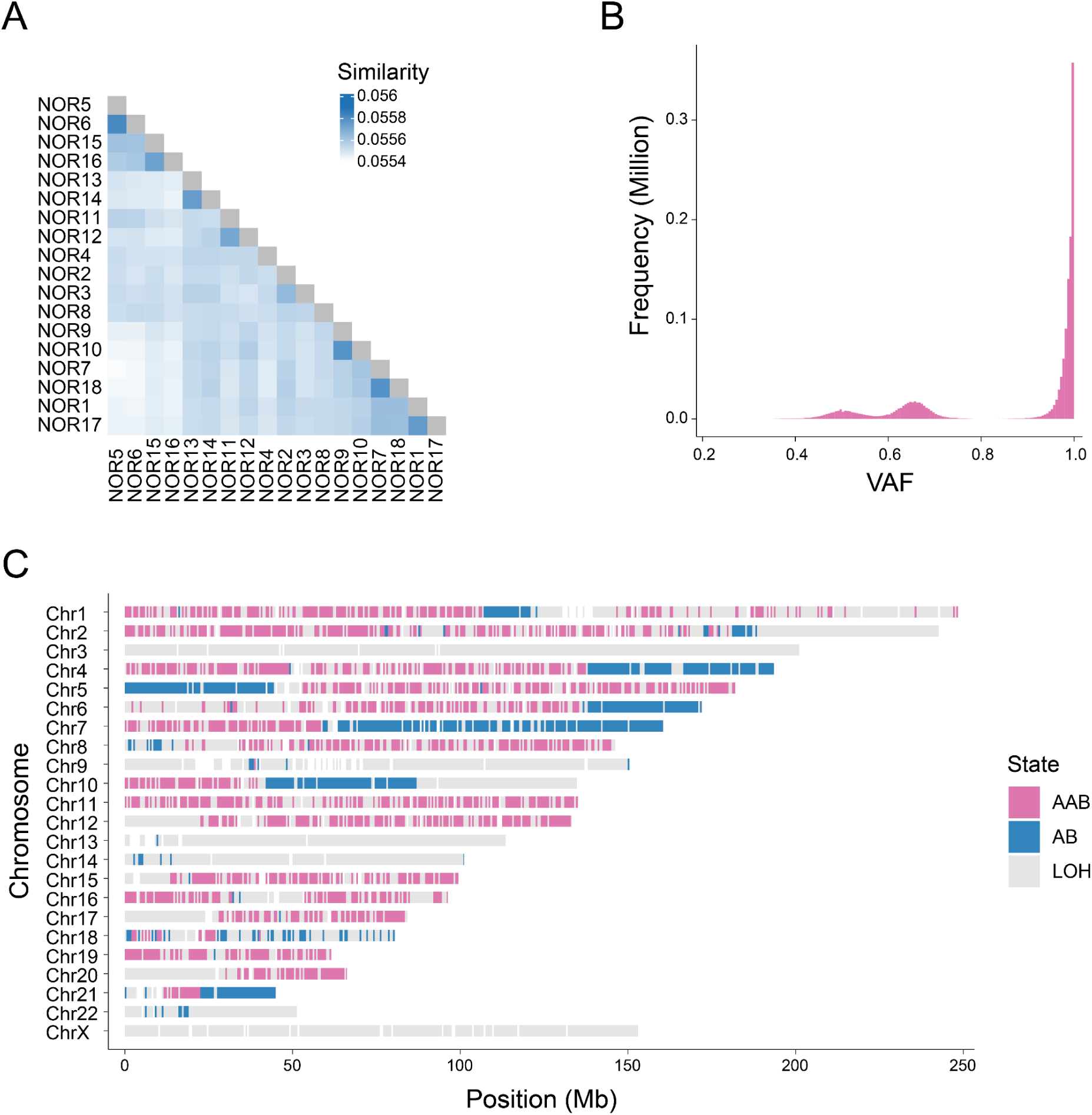
Analyses supporting chromosomal assignment of the 18 NORs. **(A)** Pairwise sequence similarity between NORs calculated using the Mash-based k-mer distance method. NORs were clustered on the similarity matrix by agglomerative hierarchical clustering (Ward’s minimum-variance method, Euclidean distance) and displayed as a heatmap. Self-similarity is not plotted. **(B)** Density plot of variant allele frequencies (VAFs) of SNPs determined using the ONT reads. **(C)** Genome-wide loss of heterozygosity (LOH) profile of the K562 genome, calculated based on VAF type frequency in non-overlapping 500-kb windows.

**fig. S7.**
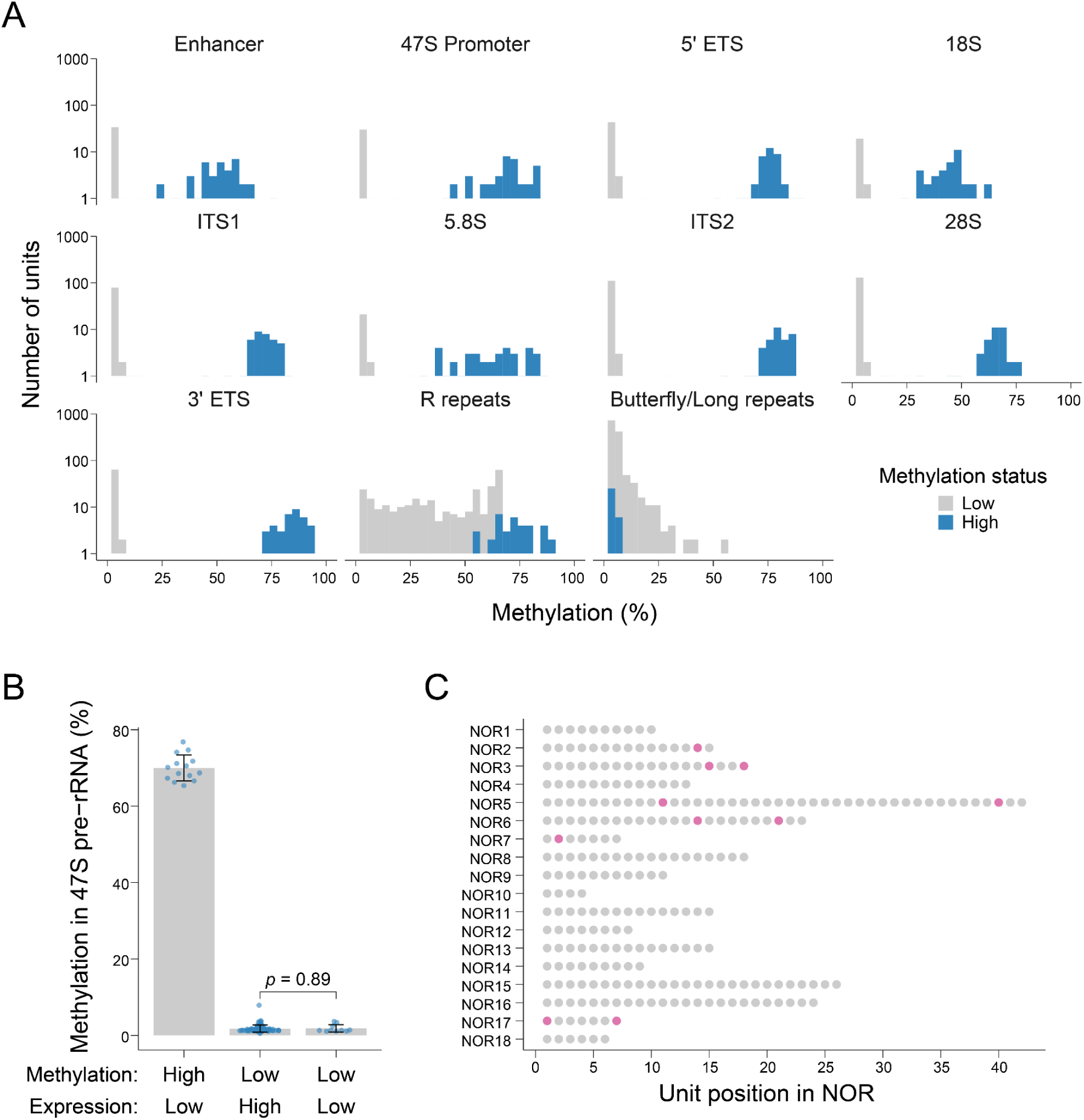
5mC methylation profiles of rDNA units. **(A)** Histograms showing DNA methylation levels across major rDNA features. Clear binary profiles are observed for most features, except for R repeats and Butterfly/Long repeats. **(B)** Bar plot comparing 47S methylation status in units marked by unique barcodes. Units are grouped as “high-methylation; low-expression”, “low-methylation; high-expression”, and “low-methylation; low-expression”. *P*-value by Welch Two Sample t-test. **(C)** Positions of the “low-methylation; low-expression” units marked by unique barcodes. The 271 full rDNA units are shown.

### Supplementary Data

**data S1**. Consensus sequences of the 18 NORs in fasta format

**data S2**. Sequence alignment of all 290 rDNA units against the reference (KY962518.1)

**data S3**. Methylation profiles of individual rDNA units

**table S1**. Statistics of ONT reads and Hifiasm outputs

**table S2**. Number of supporting reads for flank sequences

**table S3**. Truncation status of the terminal units

**table S4**. rDNA structural variation in K562, HG002, and CHM13 genomes

**table S5**. List of pegRNAs used in this study

**table S6**. List of oligos used in this study

