## Supplementary material for "Saturation barcoding of ribosomal DNA resolves the nucleolar organizer regions of the human genome": dataS2_Sequence_alignment_of_all_290_rDNA_units_against_the_reference.pdf

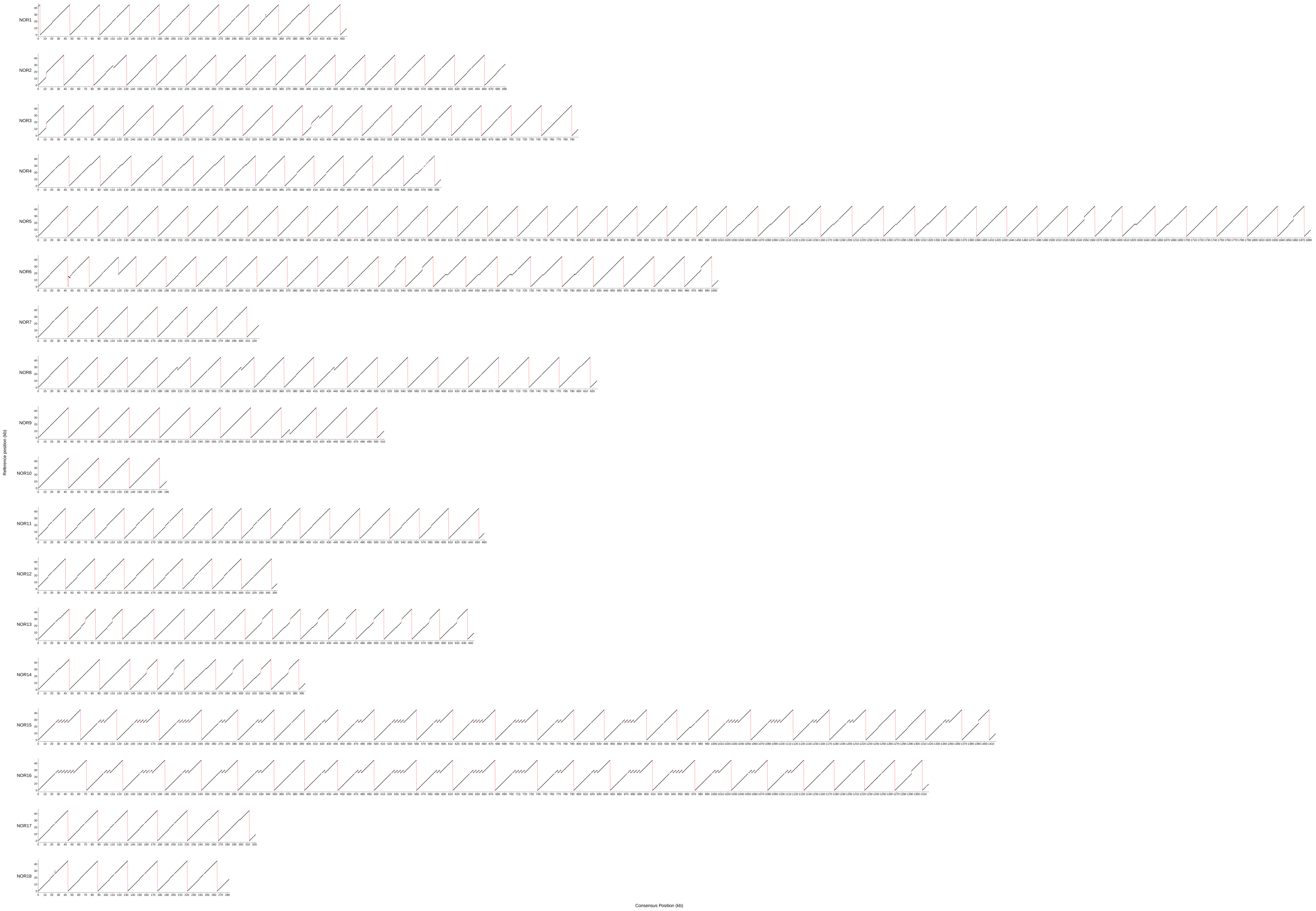
NOR1
NOR2
NOR3
NOR4
NOR5
NOR6
NOR7
NOR8
NOR9
NOR10
NOR11
NOR12
NOR13
NOR14
NOR15
NOR16
NOR17
NOR18
Reference position (kb)
Consensus Position (kb)
