## Supplementary material for "Saturation barcoding of ribosomal DNA resolves the nucleolar organizer regions of the human genome": dataS3_Methylation_profiles_of_individual_rDNA_units.pdf

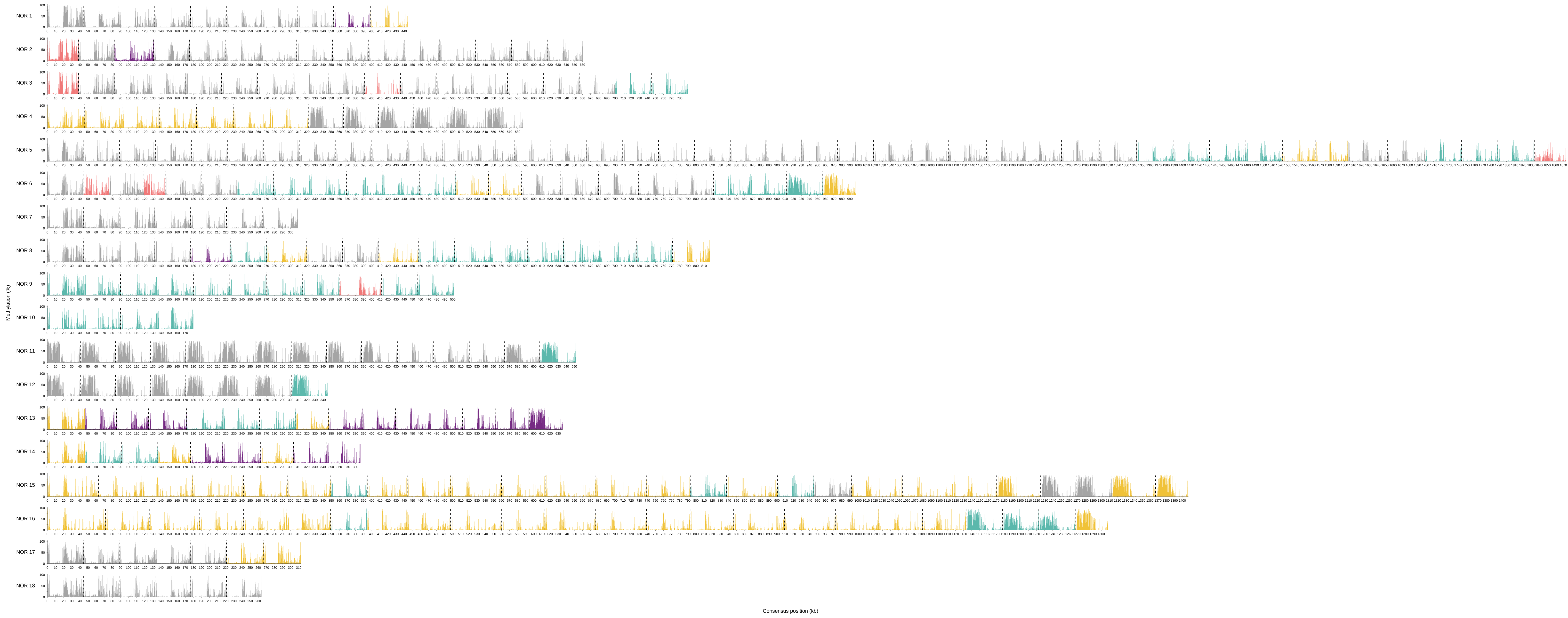

NOR 1
NOR 2
NOR 3
NOR 4
NOR 5
NOR 6
NOR 7
NOR 8
NOR 9
NOR 10
NOR 11
NOR 12
NOR 13
NOR 14
NOR 15
NOR 16
NOR 17
NOR 18
Methylation (%)
Consensus position (kb)
